# Tracking lexical and semantic prediction error underlying the N400 using artificial neural network models of sentence processing

**DOI:** 10.1101/2022.11.14.516396

**Authors:** Alessandro Lopopolo, Milena Rabovsky

## Abstract

Recent research has shown that the internal dynamics of an artificial neural network model of sentence comprehension displayed a similar pattern to the amplitude of the N400 in several conditions known to modulate this event-related potential. These results led Rabovsky, Hansen, and McClelland (2018) to suggest that the N400 might reflect change in an implicit predictive representation of meaning corresponding to semantic prediction error. This explanation stands as an alternative to the hypothesis that the N400 reflects lexical prediction error as estimated by word surprisal (Frank, Otten, Galli, & Vigliocco, 2015). In the present study, we directly model the amplitude of the N400 elicited during naturalistic sentence processing by using as predictor the update of the distributed representation of sentence meaning generated by a Sentence Gestalt model (McClelland, St. John, & Taraban, 1989) trained on a large-scale text corpus. This enables a quantitative prediction of N400 amplitudes based on a cognitively motivated model, as well as quantitative comparison of this model to alternative models of the N400. Specifically, we compare the update measure from the SG model to surprisal estimated by a comparable language model trained on next word prediction. Our results suggest that both Sentence Gestalt update and surprisal predict aspects of N400 amplitudes. Thus, we argue that N400 amplitudes might reflect two distinct but probably closely related sub-processes that contribute to the processing of a sentence.

## 1 Introduction

Understanding a sentence requires extracting semantic information about the events or states that it describes. The way this is achieved by the human brain, and more specifically the nature of the processes involved is a central question for the study of the neurobiology of language. The identification of several language-related neural correlates – such as the N400 component of the event-related potential – has allowed scientists over the last decades to have a more direct view of the neurocognition of sentence processing. Nonetheless, despite the mounting number of studies and data collected, the nature of these processes and the computations signaled by their neural correlates is still widely debated.

In this paper, we advance and test the hypothesis that the N400 event-related potential (ERP) component signals the online update of the brain’s implicit predictive representation of meaning during sentence processing, and compare it with the hypothesis that instead, it reflects lexical prediction error. We do so by mapping the internal dynamics of an artificial neural network trained on sentence comprehension onto electroencephalographic data collected during sentence reading and by comparing the results to the effect of lexical surprisal on the same data. Our cognitively motivated model consists of a re-implementation and scaling up of the Sentence Gestalt (SG) model (McClelland et al., 1989) trained on a corpus of naturalistic texts. The SG model has already been used by (Rabovsky et al., 2018) to simulate a large collection of experimental conditions modulating the N400 ERP component.

The N400 ERP component is a well-established correlate of meaning processing in the brain (Kutas & Federmeier, 2011), which offers a window into the neural processes supporting sentence comprehension. Understanding the computational processes underlying it can help us understand language and meaning processing. The N400 is a negative deflection at centroparietal electrode sites peaking around 400 ms after the onset of a word or another potentially meaningful stimulus. Its amplitude has been shown to be affected by a wide variety of linguistic variables. The N400 effect was first discovered as a larger negativity for incongruent sentence continuations (such as e.g., “I take my coffee with cream and dog”) as compared to congruent continuations (Kutas & Hillyard, 1980). In addition, N400 amplitudes tend to decrease over the course of a sentence (van Petten & Kutas, 1990). Smaller amplitudes are observed for targets after semantically similar or related as compared to unrelated primes and for repeated words as compared to a first presentation (Bentin, McCarthy, & Wood, 1985). It has been shown that the amplitude of the N400 is sensitive to the probabilistic properties of the word in isolation (e.g, its frequency; e.g., Rabovsky, Álvarez, Hohlfeld, and Sommer (2008)) and in relation to its context of utterance (e.g., its cloze probability or surprisal; (Kutas & Hillyard, 1984; van Petten & Kutas, 1990; Parviz, Johnson, Johnson, & Brock, 2011; Frank et al., 2015). Various theories propose for example that the N400 reflects, among others, lexical-semantic access to individual words or semantic integration processes at the sentence level (Kutas & Federmeier, 2011). However, in most studies, it is difficult to unequivocally decide whether reduced N400 amplitudes reflect facilitated lexical access due to prediction/pre-activation of upcoming input (Kutas & Federmeier, 2000; Lau, Phillips, & Poeppel, 2008), or whether reduced N400 amplitudes reflect facilitated semantic integration of an already retrieved word because this incoming word better fits the preceding context (C. M. Brown & Hagoort, 1993). There is also a debate revolving around the involvement of this component in predictive processes. Even though the idea of predictive preactivation underlying N400 amplitude reductions has been traditionally linked to the idea of facilitated lexical access rather than facilitated sentence level integration (Kutas & Federmeier, 2011), the issue of whether the N400 reflects predictive processing and preactivation is actually independent of the question of at which level (word or sentence level) the processes operate (Rabovsky et al., 2018). In general, despite the large body of studies conducted on the N400 and the hypothesis that its amplitude modulations are related to meaning processing, an adequate neurally-mechanistic account of the relation between the N400 and language processing with its theorized functional sub-processes is still actively debated.

Recent developments in artificial intelligence have enabled the use of computational models as explicit hypotheses for understanding cognitive processes, including language processing and its neural correlates such as the N400 ERP component. In the last decade, several studies have tried to explain the computations signaled by this component by comparing it to the performance or mechanisms implemented by various types of computational language models. These models can be ascribed to two broadly defined categories, (neuro)cognitively motivated small-scale models linking the N400 to internal processes in the models (Brouwer, Delogu, Venhuizen, & Crocker, 2021; Fitz & Chang, 2019; Rabovsky et al., 2018), and large-scale language models trained on next word prediction in naturalistic language corpora (Merkx & Frank, 2020; Michaelov & Bergen, 2020).

### 1.1 Next word prediction language models and the N400

Using next word prediction language processing models, Aurnhammer and Frank (2019); Merkx and Frank (2020); Michaelov and Bergen (2020) have shown that the N400 amplitude is significantly influenced by word-level surprisal – a probabilistic measure that quantifies how unexpected a word is given its context. More recently Michaelov, Bergen, and Coulson (2022) used surprisal estimated by state-of-the-art language models (T. B. Brown et al., 2020) and showed that it can account for many human N400 responses, but it overpredicts in some cases and underpredicts in others, such as event structure, and morphosyntactic anomalies. Additionally, a set of neuroimaging experiments provide evidence of the relation between surprisal and processing in cortical regions responsible for language processing (Willems, Frank, Nijhof, Hagoort, & van den Bosch, 2016; Lopopolo, Frank, van den Bosch, & Willems, 2017). These findings are generally taken to indicate that language processing is supported by predictive processes, reflected by the amplitude of the N400 and, by the activity in a large portion of the perisylvian and temporal cortex.

Importantly, surprisal can be estimated by a variety of language models, and thus can be implemented in different ways. These include n-gram models (also known as Markov models), recurrent neural networks (RNN’s) as well as transformer models (Jurafsky & Martin, 2009). These models can be referred to as next-word prediction language models since they are trained on predicting the next word in a sequence. In general, language models are often used to test neuro-cognitive theories of predictive processing, which posit that the brain constantly updates its expectations regarding incoming stimuli (including, but not limited to lexical units) as a function of previous inputs and conditional probabilities learned from the environment (Bar, 2011; Bubic, von Cramon, & Schubotz, 2010; K. Friston & Kiebel, 2009).

An advantage of these large-scale natural language processing (NLP) models is that they are trained on large linguistic datasets, approximating human language exposure so that they can be linked directly to empirical N400 data. However, the measure used to predict N400 amplitudes, surprisal, is an output measure of the models that can be computed in many different ways. Thus, it does not directly speak to the internal cognitive processes and neural activation dynamics underlying N400 amplitudes, which is our main interest here. Surprisal, as obtained by these language models, is a measure of the performance of the model on the task it is asked to perform – and not a measure of the internal mechanisms aimed at performing it – therefore offering a “computational” level explanation of the electrophysiological correlate in terms of Marr’s levels of analysis (Marr, 1982). Generally speaking, Marr’s computational level refers to the goal of the process carried out by a system (artificial or biological). The algorithmic level, on the other hand, refers to the the type of representations and operations involved in the process.

### 1.2 Cognitively motivated models of the N400

Cognitively motivated computational models of language comprehension that have been used to model the N400 (Laszlo & Plaut, 2012; Cheyette & Plaut, 2017; Brouwer, Crocker, Venhuizen, & Hoeks, 2017; Brouwer et al., 2021; Fitz & Chang, 2019; Rabovsky & McRae, 2014; Rabovsky et al., 2018; Rabovsky & McClelland, 2020; Rabovsky, 2020) link N400 amplitudes to internal hidden layer activation processes and dynamics. This provides an explicit and mechanistic explanation of the (neuro)cognitive processes giving rise to the N400. In other words, the link established by (neuro)cognitively motivated small-scale models between their internal processes and the N400 might instead offer an “algorithmic” (and partly “implementational”) explanation of the underlying processes.

In the first study that attempted to link neuro-cognitively motivated computational models with insights from electrophysiological studies of language processing, Laszlo and Plaut (2012) proposed that the N400 corresponds to the magnitude of the activation of a semantic representation arising from the dynamic interaction between excitatory and inhibitory connections between layers of a connectionist model of visual word recognition, and successfully simulated influences of orthographic neighborhood on N400 amplitudes. On the other hand, Rabovsky and McRae (2014) proposed that N400 amplitudes reflect an implicit prediction error at the level of meaning, which they simulated as the network error in a feature-based attractor model of word meaning. Using this model, they simulated seven empirical N400 effects, namely influences of word frequency, semantic richness, repetition, orthographic neighborhood, semantic priming, as well as interactions of repetition with both word frequency and semantic richness. Subsequently, Cheyette and Plaut (2017) addressed the same range of N400 effects as Rabovsky and McRae (2014) using a model building on Laszlo and Plaut (2012)’ proposal. The different models are not necessarily incompatible though and could be seen as complementary as the models by Laszlo and Plaut (2012) and Cheyette and Plaut (2017) address primarily the implementational level while the model by Rabovsky and McRae (2014) targets the algorithmic level of analysis according to Marr’s classification. Both models focused on N400 effects observed during the processing of single words and word pairs.

Among the small-scale models addressing N400 effects during sentence processing, Brouwer et al. (2017, 2021) tested the hypothesis that the N400 component reflects the retrieval of word meaning from semantic memory, and the P600 component indexes the integration of this meaning into the unfolding utterance interpretation. They did so by implementing the two processes – retrieval and integration – in two separate modules of a recurrent neural network-based model trained on sentence comprehension. Patterns of ERP amplitudes were compared to the internal activity changes of the retrieval and integration modules. Specifically, they monitored the dynamics of the model’s modules during the processing of reversal anomalies. Reversal anomalies are sentences such as “Every morning at breakfast, the eggs would only eat…”, where N400 amplitudes are small despite the semantic incongruity, while amplitudes of the subsequent P600 component are increased. They observed that, in this situation, the amplitude of the N400 displays similar behavior with the activity change of the retrieval module (i.e., both were small), therefore leading to their proposal that the N400 signals lexical-semantic retrieval mechanisms. On the other hand, the P600 was instead observed behaving similarly to the activity change of the integration module in the sense that both were large (see Section 4 for further discussion).

Rabovsky et al. (2018) proposed an explanation of the N400 ERP component in terms of update of an implicit predictive representation of meaning as captured by the change of the inner activation states of the Sentence Gestalt (SG) model, a connectionist model of predictive language processing that maps a sentence to its corresponding event (McClelland et al., 1989). At every given moment during sentence processing, this representation not only contains information provided by the words presented so far, but also an approximation of all features of the sentence meaning based on the statistical regularities in the model’s environment internalized in its connection weights. Rabovsky et al. (2018) showed that the SG model update (referred to as Semantic Update, or SU for short) simulates 16 distinct N400 effects obtained in empirical research. These included the influences of semantic congruity, cloze probability, word position in the sentence, reversal anomalies, semantic and associative priming, categorically related incongruities, lexical frequency, repetition, and interactions between repetition and semantic congruity. These results foster the idea that N400 amplitudes reflect surprise at the level of meaning, defined as the change in the probability distribution over semantic features in an integrated representation of meaning occasioned by the arrival of each successive constituent of a sentence. This surprise at the level of meaning has also been shown to correspond to a learning signal driving adaptation and learning in the SG model. Specifically, because at any given point in sentence processing, the model attempts to predict all aspects of meaning of the described event, the change in SG activation induced by each new incoming word corresponds to the prediction error contained in the previous SG representation. Based on the idea that prediction errors drive learning, Rabovsky et al. (2018) used the difference in SG activation between the current and the next word as a learning signal to adjust connection weights in the model (simulation 16). This was taken to suggest that N400 amplitudes reflect an implicit error based learning signal during language comprehension (Rabovsky et al., 2018). Thus, the model by Rabovsky et al. (2018) refined the notion of an implicit prediction error at the level meaning proposed by Rabovsky and McRae (2014) by extending it to the sentence level, directly linking it to neural activation, and implementing it dynamically as a change in an implicit predictive representation of meaning.

Fitz and Chang (2019) extended the perspective that N400 amplitudes reflect a learning signal also to the P600. Specifically, they proposed that both the N400 and P600 arise as side effects of an error-based learning mechanism that explains linguistic adaptation and language learning (Gehring, Goss, Coles, Meyer, & Donchin, 1993; Holroyd & Coles, 2002). Their model is based on Chang’s Dual-path model – a connectionist model of language acquisition and sentence production (Chang, 2002). The model decouples sequence and meaning processing in two distinct pathways, which – during training – learn syntax and semantic regularities. Their theory is instantiated by observing how the model’s processing error simulates data from several studies on the N400 (amplitude modulation by expectancy, contextual constraint, and sentence position), and on the P600 (agreement, tense, word category, sub-categorization and garden-path sentences).

These cognitively motivated models proposed to account for N400 amplitudes have as of yet been only trained on small artificial language corpora. This has some advantages with respect to the models’ transparency but also entails important limitations. Specifically, the models’ N400 correlate – its internal activation dynamics – can be related to empirical N400 data only in a qualitative way and only concerning specific experimental manipulations. For instance, even though Rabovsky et al. (2018) showed that the SG model covers a wide range of distinct N400 effects, including ones that could not be accounted for by lexical surprisal, quantitative assessment of the relation between its activity and the amplitude of the N400 measured in empirical studies as well as quantitative model comparison was impossible. This is because it was trained only on a small synthetic language and therefore it could not be presented with the same stimuli presented in empirical experiments. For this reason, the relation between the model’s N400 correlate and empirical N400 data remained somewhat abstract.

### 1.3 The present study

In the present study we overcome this limitation and quantitatively investigate the hypothesis that N400 amplitudes might reflect predictive processes at the level of sentence meaning based on our cognitively motivated SG model trained on a large scale corpus. Moreover, we quantitatively compare this hypothesis with the alternative hypothesis that they instead signal prediction error at the word level by implementing a next word prediction LM. From the SG model we estimate the change over successive words of its internal representations of the meaning of the sentence, otherwise known as Semantic Update (SU.SGM). Whereas, the LM, trained on next word prediction, is employed to estimate word-level surprisal, a measure that has already been shown to correlate with the N400 (e.g., Frank et al., 2015). Using SU.SGM based on the internal dynamics of the SG model implies that N400 amplitudes reflect predictive processes at the level of sentence meaning. Whereas, using surprisal derived from a next word prediction language model (LM) can be seen as implementing the idea that the N400 reflects lexical prediction and prediction error (but see section 4.3 in the discussion section for an alternative interpretation based on Levy, 2008).

In the analyses conducted in the present study, we use these measures to directly predict the amplitude of the N400 generated during sentence processing. Moreover, in order to make the quantitative comparison between the effects of SU.SGM and surprisal meaningful and fair, we compute this latter measure from a language model having an architecture as similar as possible to the SG model and that has been trained on the same linguistic material. This ensures, in our view, that the models’ different performances with regard to the EEG data are not due to differences in training data or (as far as possible) basic architectural features, but that these differences are ascribable to differences in training task (next word prediction versus predictive meaning comprehension) and to the comparison between a performance-oriented measure (surprisal) and an internal-processing one (SU.SGM). To further examine the impact of the training task on the internal representations generated by the models during sentence processing, we also explore the relationship between the N400 and the update of the internal representations generated by the next word prediction language model (SU.LM, Supplementary MaterialB).

The findings presented in this paper show that both SU.SGM and surprisal have a significant effect on the amplitude of the N400 and the temporal pattern of EEG activity from 300 to 500 ms after word onset. These effects persist even when accounting for surprisal and SU, respectively, indicating that these two measures potentially address distinct aspects or processes within the broader range of activity triggered by sentence comprehension. In comparison to lexical surprisal, our measure of Semantic Update in the SG model exhibits a more prolonged temporal resemblance to brain activity, extending beyond the conventional N400 time-frame.

### 1.4 The Sentence Gestalt model

The **Sentence Gestalt (SG) model** is a model of language comprehension which maps sentences to a description of its meaning approximated by a list of arguments encompassing the action, the various participants (e.g., agent and patient) as well as information concerning, for instance, the time, location, and the manner of the situations or events conveyed by the speaker (McClelland et al., 1989). For instance, when processing the sentence “the boy opened the door slowly”, the **task** of the SG model is to recognize that *opened* is the action, and that *the boy* and *door* are its agent and patient respectively and that *slowly* is a modifier specifying the way in which the event takes place. In order to process such a sentence, the model implements a mechanism consisting in building internal representations (here referred to as Sentence Gestalts) that can be used as a basis to respond to probes regarding the meaning of the sentence approximated by the argument structure of the utterance in terms of role filler pairs. The model implements a theory of sentence processing centered around the computation of an internal representation of meaning informed and constrained by the sentence uttered by the speaker.

The model is probabilistic in the sense that it can be conceptualized as approximating a function computing a conditional probability distribution between role-filler pairs, representing the arguments of the sentence’s event, and a string of words composing the sentence itself: *P*(*< r, f > |w*_1:*n*_), where *r* refers to the role (e.g., *agent*) and *f* to the filler (e.g., *boy*) of an argument, and *w*_1:*n*_ is the sequence of the first *n* words of a sentence. For instance, given the sequence *the, boy, opened*, the model learns the conditional probability of the role-filler pair *< agent, boy >*. This is done on the basis of the statistical properties of the training environment and by using probes. Given a sentence and a probe, the training environment specifies a probability distribution over the possible answers to the probe and the goal of processing is to form a representation that allows the model to match this probability distribution. Moreover, the SG model is predictive in light of its training regime, which forces it to estimate role-filler pairs representing arguments of the whole sentence event structure word-by-word, even before the actual lexical items that correspond to the probed argument fillers are available. The model is therefore expected to attempt to use the representation to anticipate the expected responses to these probes. Section 2.2 below contains the details regarding the training procedure adopted in this study. Finally, the predictive and probabilistic processing at the level of meaning is internally represented in the model by the hidden Sentence Gestalt representations. These representations are implicit in that they represent sentence meaning using features that do not correspond to symbolic features, either semantic or grammatical. That makes them formally comparable to semantic representations obtained from vector-space models such as the ones produced by distributional semantics and deep neural network word embeddings.

As stated in Section 1.3, the SG model is introduced as an alternative to a **next word prediction language models (LMs)** in light of aspects of the theory of language processing that they implement. Despite the fact that both models are predictive, LMs implement a theory of prediction at the level of lexical items. In contrast, as seen above, SG models are trained to predict sentence meanings from sequences of words. In line with this, both models approximate functions estimating a conditional probability distribution. If the SG model estimates conditional probabilities between lexical cues and role-filler pairs of event arguments, an LM instead estimates such probabilities between sequences of words and their following word. Given a sequence of words of length *n*, an LM assigns a probability *P*(*w_n_*_+1_*|w*_1:*n*_) to the next word *w_n_*_+1_.

## 2 Materials and methods

### 2.1 The implementation

The **SG model** is constituted of two components: an update network (encoder) and a query network (decoder), as described in Figure 1. The update network sequentially processes each incoming word to update activation of the Sentence Gestalt layer, which represents the meaning of the sentence after the presentation of each word as a function of its previous activation and the activation induced by the new incoming word. The query network, instead, extracts information concerning the event described by the sentence from the activation of the Sentence Gestalt layer and it is primarily used for training.

**Fig. 1:**
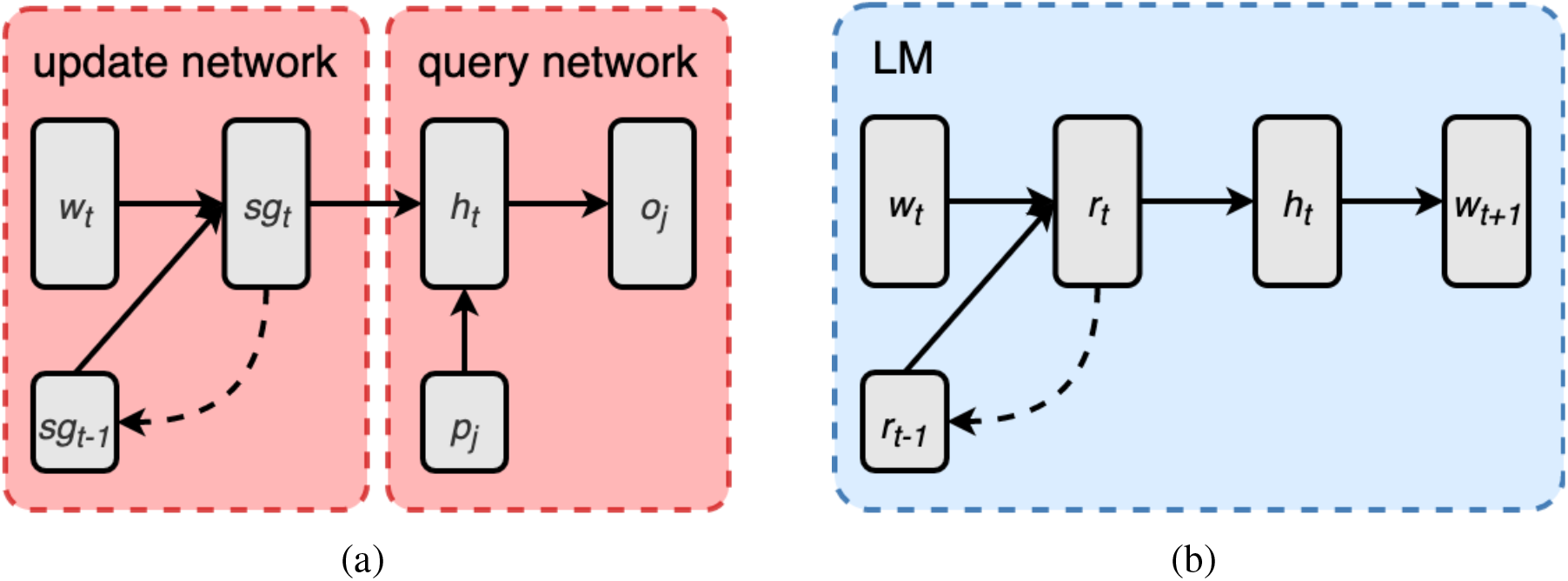
The architecture of the SG model (a) and the LM (b).

The **update network** of the SG model is composed of an input layer, which generates a learned vectorial representation 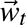 for each input word of the incoming sentence, and a recurrent layer implemented as a long short-term memory (LSTM) unit generating a Sentence Gestalt representation 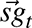 as a function of the current input word 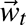 and its previous Sentence Gestalt representation 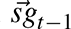 (Hochreiter & Schmidhuber, 1997). LSTM have the advantage of being better at processing long and complex sentences compared to traditional recurrent layers, and being still simpler in structure and number of parameters compared to even more performative types of deep learning components (e.g. Transformers). The update network is essentially a recurrent neural network encoding a string of words as a function of the current presented word and the words preceding it. The **query network** is composed by a hidden layer 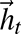 which combines the Sentence Gestalt vector 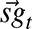 and a probe vector 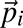. The output 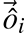 of the query network is generated from the hidden state 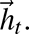

The **task** the SG model is asked to perform is to map a sentence to its corresponding situation or event, defined as a list of role-filler pairs representing an action or state, its participants (e.g. agent, patient, recipient), and eventual modifiers. A sentence is defined as a sequence of words, each represented as an integer *i_t_*, defined on a lexicon associating a unique index to every word. Figure 2 exemplifies the mapping from words to event performed, word-by-word, by the SG model. The left column (a) displays a toy example of the role-probing procedure where given the sentence *the boy opened the door*, the model tries to identify the role, for instance, of the word *boy*. The sentence is presented word-by-word. The model is first presented only with the words *the boy*. At this stage, after receiving only the first words of a sentence (a.i), the model generates a list of possible roles for the filler word *boy* (AGENT, PATIENT, BENEFICIARY), which are updated as new information, in the form of subsequent words, is presented by the input (a.ii). On the right (b) we show an example of the filler-probing procedure where given the sentence *the boy opened the door*, the model tries to identify the filler of the *patient* role. The sentence is presented word-by-word. The model is first presented only with the words *the boy* (b.i), leading to a list of wrong predicted patient fillers. As soon as the model is presented with word *opened* (b.ii), its prediction is adjusted to produce a list of potentially correct fillers. When the model is presented with the whole sentence (b.iii), it converges on the correct patient of the described event: *door*.

**Fig. 2:**
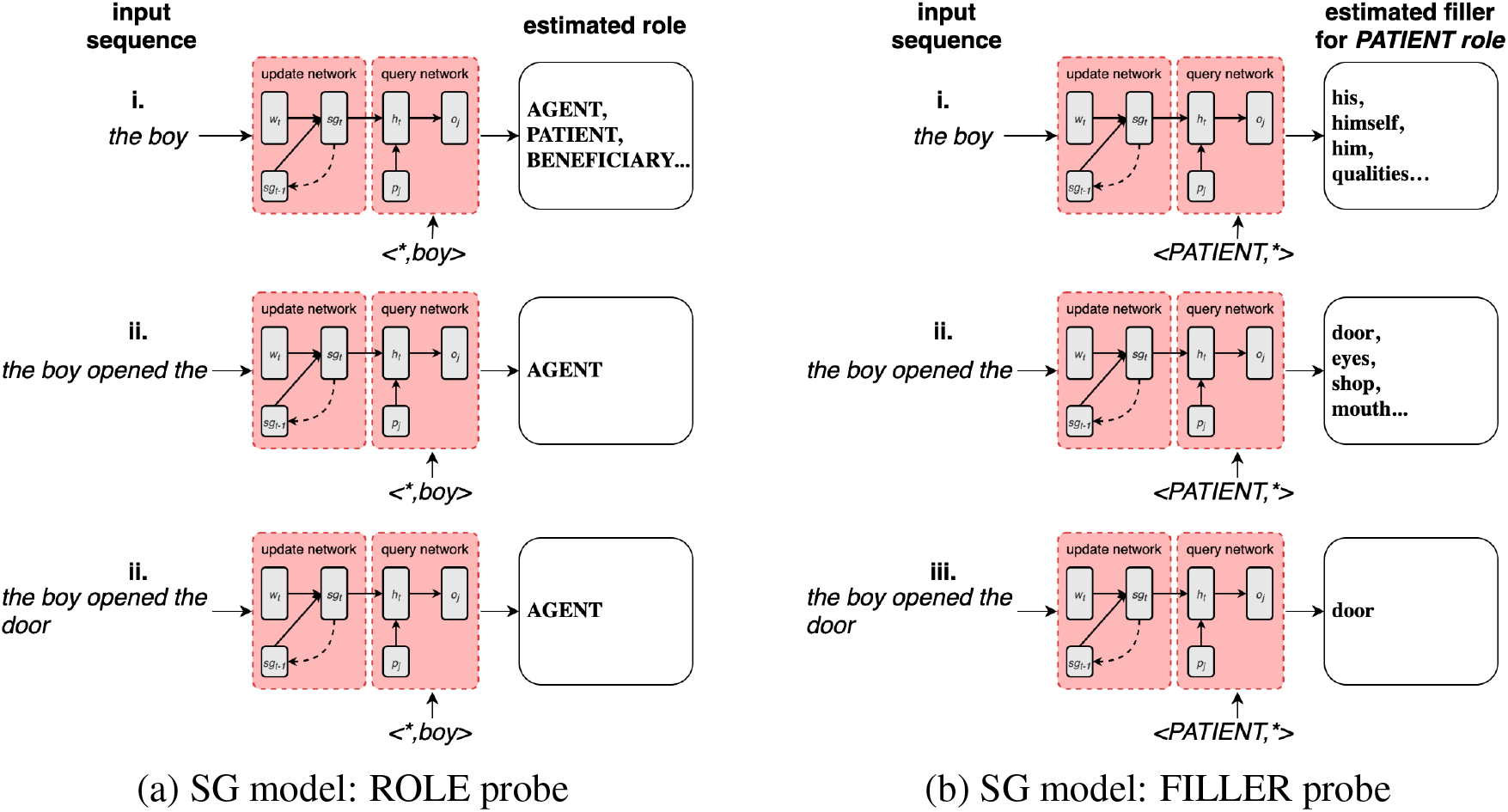
Example of how the SG model processes the sentence the boy opened the door. The model is trained on identifying the roles (a) and fillers (b) constituting the event described in the sentence. In both cases the sentence is presented word-by-word. The model is interrogated on all the role-filler pairs of the event described by the sentence. **(a)** ROLE probing: after receiving only the first words of a sentence (a.i), the model generates a list of possible roles for filler boy (AGENT, PATIENT, BENEFICIARY), which are updated as new information, in the form of subsequent words, is presented by the input (a.ii). **(b)** similarly for FILLER probing: when the model is first presented only with the words the boy (b.i), this leads to a list of wrong predicted patient fillers. Subsequently (b.ii), the model is presented with word opened, causing the prediction of potentially correct patient fillers. When the model is presented with the whole sentence (b.iii), it converges on the correct patient of the described event: door.

As shown in Figure 3.a, the event consists of a set of role-filler vectors 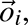 each of which consists of the concatenation of the feature representation of a word (the filler) and a one-hot vector of the role of that word in the context of the event described by the sentence. Therefore, the sentence “the boy opened the door slowly” will consist of a sequence of 6 one-hot word representation vectors. Its event contains 4 role-filler combinations representing each role of the event (agent, action, patient, manner) with its corresponding concept filler (boy, open, door, slowly).

**Fig. 3:**
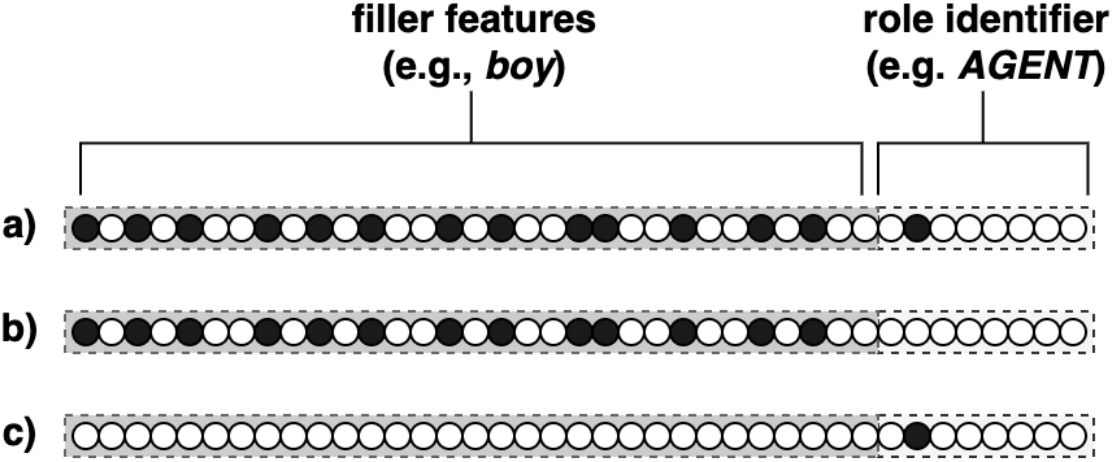
The role-filler vector 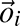 (a), and its corresponding two types of probes 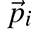 (b) and (c). The left hand-side of the vectors correspond to the embedding representation of the filler concept, whereas the right hand-side to the one-hot representation of the thematic role played by the filler. When probing for the thematic role, probe (b) is presented. When probing for the filler instead, probe (c) is presented. In both cases the SG model is expected to produce the full role-filler vector (a).

During **training** the model is presented with sentences such as “the boy opened the door”, word-by-word, and their corresponding role-filler list (*< boy, Agent >*, *< opened, Action >*, *< door, Patient >*). The model is presented with the following sentence-specific probes: (*< boy, * >*, *< opened, * >*, *< door, * >*, *< *, Agent >*, *< *, Action >*, *< *, Patient >*). In the present formulation of the model, only probes corresponding to actual roles and fillers describing the sentence are presented; for instance, the model will not be probed for a role describing the temporal modifier of the event, since such modifier is not present in this specific training sentence. All probes are presented one by one after the presentation of each word. For instance, the model sees the word “the” at the input layer and subsequently is probed with each probe listed above (for instance *< boy, * >*) and expected to generate the corresponding complete role-filler pair relative to this sentence at the output layer, in this case: *< boy, Agent >*. This is repeated for all probes and after the presentation of each lexical input. Notably, the SG model is probed even before the relevant lexical item, like “boy”, is presented to the input layer. This approach forces the model to learn a predictive function that operates at the level of sentential meaning. A probe consists of a vector 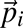 of the same size of a corresponding role-filler vector 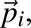 but with either the thematic role identifier zeroed (Figure 3.b) – if probing for roles –, or filler features zeroed (Figure 3.c) – if instead probing for fillers. Responding to a probe consists therefore of completing the role-filler vector. When probed with either a thematic role (e.g., agent, action, patient, location, or situation; each represented by an individual unit at the probe and output layer) or a filler, the model is expected to output the complete role-filler vector. Fillers are represented using word embeddings obtained by binarizing *Fasttext*, a computational semantic model representing 1 million words and trained on both the English Wikipedia and the Gigaword 5 corpora (Bojanowski, Grave, Joulin, & Mikolov, 2017)^1^. The discrepancies between the observed role-filler vector 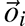 and generated output 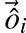 is computed using cross-entropy and is back-propagated through the entire network to adjust its connection weights in order to minimize the difference between model-generated and correct output. Binarization of the filler semantic feature representations was performed in order to allow for a probabilistic interpretation of the model generated activation of semantic feature units afforded by the cross-entropy error used during training.

Besides being very powerful and widespread tools in natural language processing (NLP), **next word prediction LMs** have been used, as mentioned in the introduction, in numerous studies of language processing in humans, in psycholinguistics and neurobiology of language (Frank et al., 2015). RNN implementations of the LMs share significant similarities in the way they treat the linguistic input and in internal architecture with the SG model. In both cases language – for instance, sentences – are treated as sequences of words which are fed to an input layer which generates a per-word vectorial representation which is then integrated with a contextual representation by the internal recurrent layer. The LM is composed of an input layer, which generates a vectorial representation 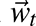 for each input word of the incoming sentence, and a recurrent layer implemented as a long short-term memory (LSTM) computing a recurrent representation 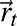 as a function of the current input word 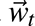 and its previous activation 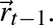 The hidden layer 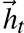 takes the recurrent representation 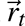 and feeds it to the output 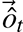 which consist of a probability distribution over the model’s lexicon representing the likelihood of the next word *w_t_*_+1_.

The structural differences between the SG and LM models are purposefully designed to align with their respective functions of lexical and sentential-semantic prediction. It is essential to emphasize that these differences are not arbitrary, but rather driven by theoretical considerations. While the models exhibit similarities in their input, recurrent, and post-recurrent feed-forward layers, they deliberately diverge in their output layers to effectively tackle their designated tasks. These intentional variations emphasize the significance of the models’ theoretical motivations and underscore the crucial distinction between lexical and sentence meaning prediction, which forms the central focus of our study.

### 2.2 Training the models

Both LM and SG model are trained on the same data, the Rollenwechsel-English (RW-eng) corpus (Sayeed, Shkadzko, & Demberg, 2018). The only crucial difference is – as mentioned above – the task the two models are called to perform: predict the next lexical item (i.e. mapping sequences to words) for the LM, or understanding and predicting the meaning of the sentence (i.e. mapping sentences to role-filler pairs) for the SG model.

The RW-eng corpus is annotated with semantic role information based on PropBank roles (Palmer, Gildea, & Kingsbury, 2005) and obtained from the output of the SENNA semantic role labeller (Collobert et al., 2011; Collobert, 2011) and the MALT syntactic parser (Nivre, 2003). Each sentence annotation consists of the list of event frames it describes. An event frame is defined by its main predicate (usually a finite verb) and its arguments. Following PropBank, RW-eng frames can contain arguments of 26 types spanning from agent, patient, benefactive, starting and end point and a series of modifiers describing the temporal, locational, causal, final and modal circumstances of the event. Therefore, the SG model in this study is trained on mapping each RW-eng sentence to its PropBank-style event structure as provided in the RW-eng corpus. For more detail on the argument structure proposed by PropBank we refer to Palmer et al. (2005). A sentence can contain multiple event frames. For instance, consider the following sentence (taken from the RW-eng corpus):

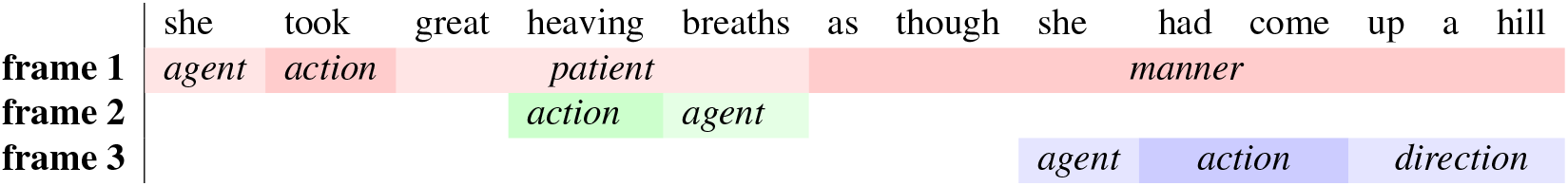

The sentence is composed by 13 lexical items and describes 3 nested events represented by 3 separate PropBank-style frames. The first, color-coded in red, contains a patient (“great heaving breaths”) and a manner modifier (“as though she had come up a hill”) which in turn are frames on their own right (color-coded in green and blue respectively). It is worth noting that the SG model is a model of language comprehension in general, and that it is trained not only on events. The training dataset includes also other types of utterances such as stative sentences, questions and commands among others. This example illustrates how the SG model is trained on naturalistic, complex, English sentences, by presenting it with one word at a time, following the sentential order, and probing each time for the role-fillers of the argument of potentially several events at the same time.

Instead, the LM was trained only on the raw sequences of words (sentences) contained in the corpus, disregarding the semantic annotation based on PropBank. When trained on the example sentence above, the model is asked to predict the simple linear sequence of lexical items. It is presented, for instance with the input sequence “…she took great heaving” and asked to match it with the next lexical item “breaths”, as per the example sentence above.

The connection weights of both models were optimized using Adamax (Kingma & Ba, 2015) with learning rate equal to 0.01 for the SG model and 0.002 for the LM (after hyperparameters optimization on 5% of the data). The whole dataset was split in mini-batches of 32 sentences each. Training was conducted for a maximum of 150 epochs on 90% of the batches, the remaining 10% was kept for validation. Only sentences having between 6 and 15 words and having a maximum of 8 frames were used for training. Sentence length and number of frames were constrained in order to limit the number of complex subordinate events and to facilitate the mini-batch training.

The size of the input layers of both model was equal to 600 and their recurrent layers size equal to 1200 (the Sentence Gestalt layer for the SG model), implemented as a 1-layer LSTM. The SG model’s probe and output layers had size 328 due to the concatenation of the 300-size binarized embedding vector, the frame number and the argument type. Instead the output layer of the LM was equal to the extension of its lexicon: 300000.

### 2.3 Model-based variables

The goal of using these two modelling approaches is to investigate the relation between brain activity (more specifically N400 amplitudes and the related EEG time-course post word onset) and the update of an internal predictive representation of sentence meaning, on one hand, and lexical prediction error, on the other. For this reason we compute two main measures from our models: Semantic Update (SU) and surprisal.

Semantic Update is the update of the SG model’s recurrent layer’s internal representations after the presentation of each word composing a sentence. It is computed as the mean absolute error between the activation of the Sentence Gestalt layer after the presentation of a word and its activation before the presentation of that word. It represents the amount of change driven by the new incoming word to the implicit predictive representation of sentence meaning. SU is computed from a trained SG model, and SU computation and training involve two separate sets of data: the stimulus sentences used in the empirical EEG study (for computing SU) and the training corpus (for training). During SU computation, the update module of the SG model reads the stimulus sentences as sequences of words encoded as 1-hot vectors. No additional input information, such as probing as in the case of training, is provided at this stage, and therefore the probes and targets are not directly used to compute the Sentence Gestalt activation and its update.

Surprisal instead is the negative log-probability of the new incoming word given the previous words: *surp*(*w_t_*)= *-log*(*P*(*w_t_|w*_1:*t*_*_-_*_1_)). It is computed from the probability distribution generated as output of the LM. Surprisal operates on lexical items, and not on the semantic representations, and is taken to represent the error in predicting such items in a sequential input.

Additionally, in Supplementary Material B in order to compare the effect of training task on the update of the internal representation of a model and to consider an algorithmic level measure from the LM as well, we computed an LM equivalent of the SG model’s SU by tracking the update of the LM’s recurrent layer’s internal representations after the presentation of each word composing a sentence. We refer to this update variable as the SU.LM in order to distinguish it from the one obtained from the SG model (SU.SGM).

### 2.4 EEG dataset

The elecrophysiological recordings of the N400 were obtained from an EEG dataset provided by Frank et al. (2015). The dataset consists of data collected from twenty-four participants (10 female, mean age 28.0 years, all right handed and native speakers of English) while they were reading sentences extracted from English narrative texts.

The stimuli consisted of 205 sentences (1931 word tokens) from the UCL corpus of reading times (Frank, Monsalve, Thompson, & Vigliocco, 2013), and originally from three little known novels. The sentences were presented in random order, word by word. About half of the sentences were paired with a yes/no comprehension question to ensure that participants read attentively. The sentence stimuli were not manipulated to control or elicit any particular linguistic phenomenon.

The sentences were presented in random order, word-by-word. Each word was presented in the center of the screen for a variable amount of time, depending on its length as number of characters. Word presentation duration equalled 190 ms plus 20 for each character in the word. Each word was followed by a 390 ms interval before appearance of the next word.

The EEG signal was recorded continuously at a rate of 500 Hz from 32 scalp sites (Easycap montage M10) and the two mastoids. Signal was band-pass filtered online between 0.01 and 35 Hz. Offline, signals were filtered between 0.05 and 25 Hz (zero phase shift, 96 dB roll-off). The N400 amplitude for each subject and word token was defined as the average scalp potential over a 300-500 ms time window after word onset at electrode sites in a centro-parietal region of interest (ROI), which corresponded to the ROI used by Frank et al (2015).

For further details regarding the stimuli see Frank et al. (2013). More detailed information regarding the EEG dataset, its stimulation paradigm, preprocessing, and the electrode positions contained in the ROI can instead be found in Frank et al. (2015).

## 3 Analyses

In this study we want to test the hypotheses that language-elicited EEG activity, specifically the amplitude of the N400 component, reflects prediction errors at the level of sentence meaning or lexical-items.

In Section 3.1 we fit a linear mixed effect model with the aim of predicting the amplitude of the N400 (average activity over a 300-500 ms time window after word onset at centro-parietal electrodes, see Supplementary Material A) as a function of the update of the semantic representation generated by the SG model during language processing (the SU.SGM) and as a function of surprisal computed from a LM. Figure 4 presents a graphical summary of our approach. Since the N400 is a negative deflection of the electrophysiological signal, the SU.SGM is multiplied by *-*1.

**Fig. 4:**
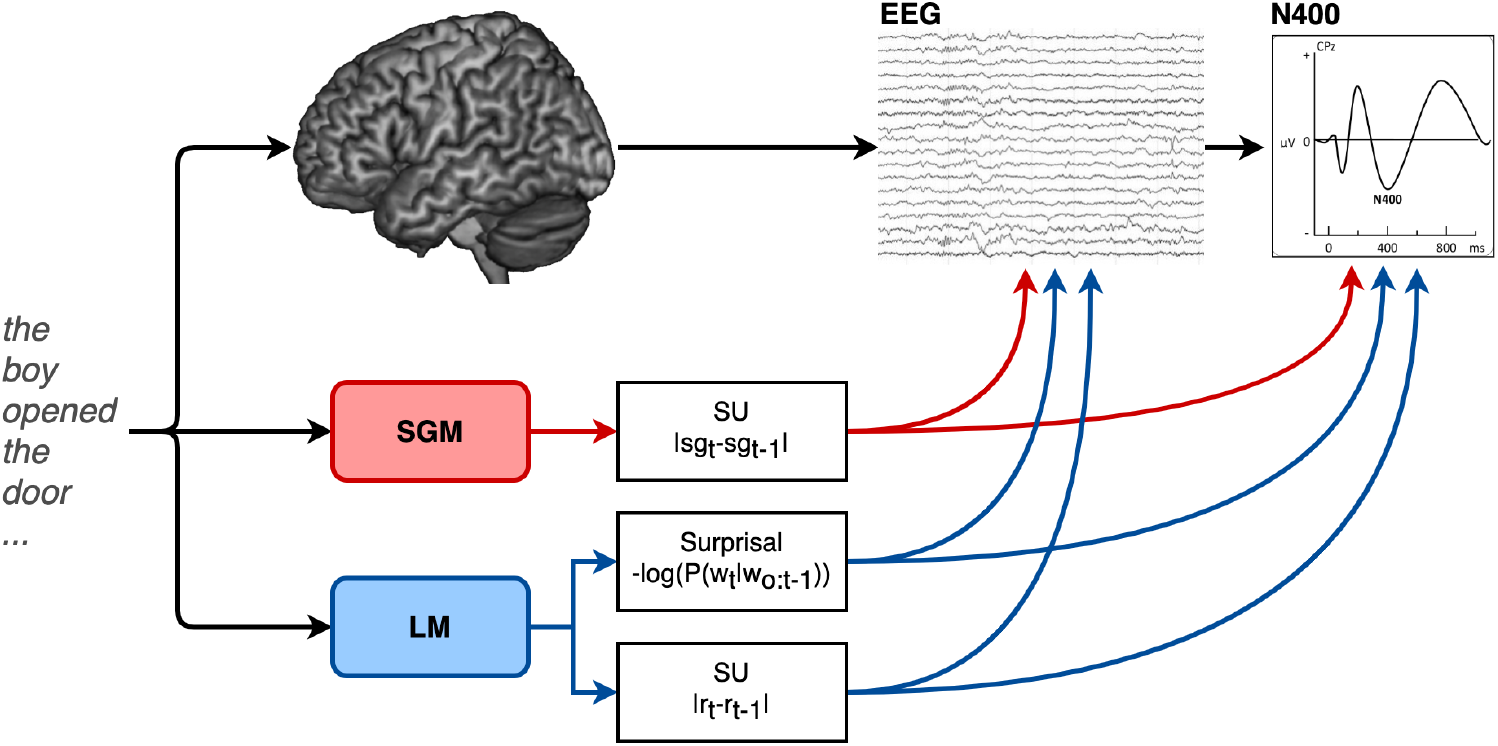
This study aims to model the amplitude of the N400 and the electrophysiological activity locked to the presentation of each word during sentence comprehension as a function of the update of an implicit predictive semantic representation generated by a SG model (SU.SGM) trained on a large scale corpus of naturalistic texts. The effect of SU.SGM is compared to the one of surprisal as computed by a LM and to the internal update of the LM (SU.LM).

In Section 3.2, we assess the contribution of the SU.SGM above and beyond the effect of surprisal, and vice versa. Since the information reflected by our predictors might go beyond the time window of the N400 most typically defined as the average per-trial activity between 300 and 500 ms post-word onset, and to investigate the specificity of the observed relationship between SU.SGM and N400 amplitudes, in Section 3.3 we predict the complete time-course of the EEG signal in order to explore whether earlier or later latencies also correlate with the SG model’s internal dynamics. These analyses are compared and contrasted to the effect of lexical surprisal on the same electrophysiological measurements.

SU.SGM has a correlation of 0.16 with surprisal.LM (see Figure 5).

**Fig. 5:**
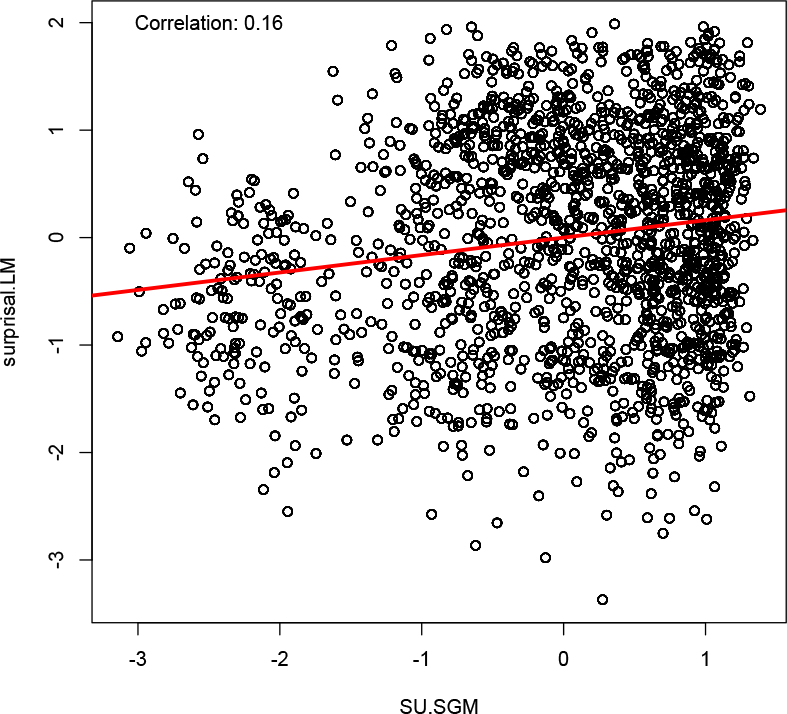
Correlation between surprisal and the SU from the SG model.

### 3.1 Predicting the N400

We predict the amplitude of the N400 ERP component amplitude obtained from Frank et al. (2015) as a function of either the update of the SG model (SU.SGM, Table 1) and word surprisal estimated by the LM (surprisal.LM, Table 2) computed over the stimulus words. All linear models include the ERP baseline (the activity of the 100 ms leading to the onset of each word). In order to avoid potential artefacts, the baseline is not subtracted directly from the dependent variable, but instead included as a variable of no interest in the models. The models are fit with per-subject random slopes and random intercepts, and per-word random intercepts. The N400 was computed as the average activity between 300 and 500 milliseconds after word onset in centro-parietal electrodes (see Supplementary Material A for a depiction of the electrode set-up and the electrodes selected for analysis, based on Frank et al. (2015)).

**Tab. 1:**
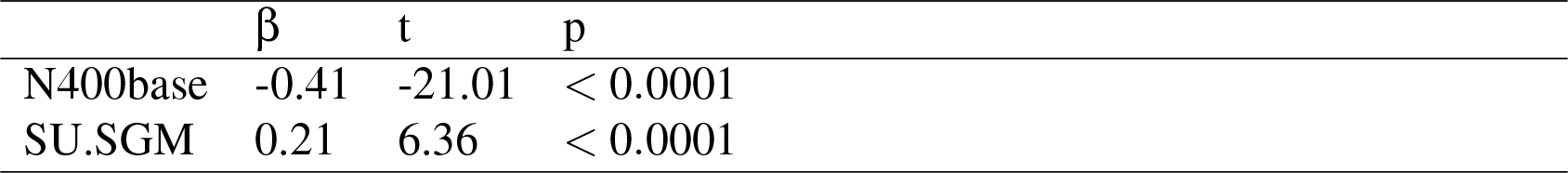
Linear mixed effect model fitted with the update of the SG model (SU.SGM) and aimed at predicting the amplitude of the N400 component.

**Tab. 2:**
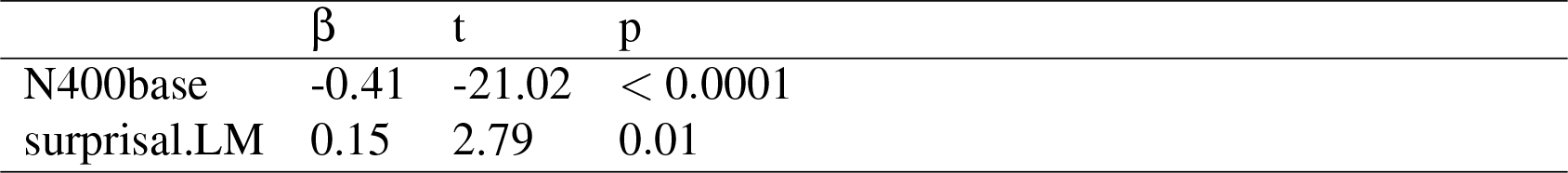
Linear mixed effect model fitted with surprisal estimated by the LM (surprisal.LM) and aimed at predicting the amplitude of the N400 component.

No other factors, such as position and frequency, were included in the current analyses as control variables. Our introduction of SU.SGM aims to provide a computationally explicit theory of the cognitive mechanism underlying the N400, rather than proposing an additional variable that influences the N400 beyond existing variables. This cognitive mechanism (semantic update), should thus be influenced by the same factors that affect the N400, including sentence position and lexical frequency. Our starting point is Rabovsky et al. (2018) which demonstrated this point by showing that numerous factors known to influence the N400, such as sentence position, lexical frequency, and semantic relatedness, influenced semantic update in the same way. Therefore, from our theoretical perspective, the influences of variables that modulate the N400 contribute to the variance we aim to explain and should not be controlled for^2^.

The results in Table 1 clearly indicate that SU.SGM significantly predicts the amplitude of the N400 (β = 0.21, *t* = 6.36, *p <* 0.0001 FDR corrected). This indicates that larger word-wise updates of the Sentence Gestalt layer representation correspond with stronger negative deviation of the ERP signal in the N400 time segment.

Table 2 shows that the surprisal.LM has a significant effect on the amplitude of the N400 (β = 0.15, *t* = 2.79, *p* = 0.01 FDR corrected).

### 3.2 Comparing surprisal and update as predictors of the N400

In this section we compare the effect of Semantic Update and surprisal as predictors of the amplitude of the N400. We first fit a linear mixed effect model predicting the amplitude of the N400 using as predictors both SU.SGM and surprisal estimated from the LM. The models contains also the N400 baseline and is fit with per-subject random slopes and random intercepts, and per-word random intercepts.

Table 3 contains the results of a linear mixed effect model fitted with SU.SGM and surprisal, showing that both variables have a significant contribution to the prediction of the amplitude of the N400 (surprisal.LM: β = 0.16, *t* = 3.02, *p* = 0.006 FDR corrected; SU.SGM: β = 0.22, *t* = 8.36, *p <* 0.0001 FDR corrected).

**Tab. 3:**
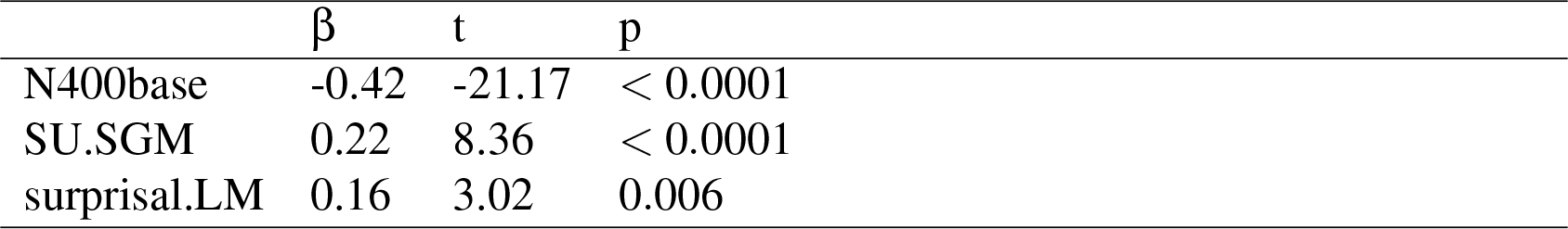
Results of a model fitted with both surprisal and SU.SGM and aimed at predicting the amplitude of the N400 component.

| | $\beta$ | t | p |
| --- | --- | --- | --- |
| N400base | -0.42 | -21.17 | $< 0.0001$ |
| SU.SGM | 0.22 | 8.36 | $< 0.0001$ |
| surprisal.LM | 0.16 | 3.02 | 0.006 |

We assessed the improvement in the prediction of the N400 by incorporating either the SU.SGM or surprisal.LM into a model fitted only with the N400 baseline. The models were fitted with per-subject random slopes and random intercepts, along with per-word random intercepts. We conducted a two log-likelihood test between the models reporting both χ^2^ and Δ*AIC*. AIC (Akaike Information Criterion) is a statistical measure used to compare the goodness-of-fit of different models while taking into account model complexity. Lower AIC values indicate better model fit. When comparing two nested models, the Δ*AIC* is the difference between the AIC values of the two models.

The results reported in Table 4 indicates that both the addition of SU.SGM and of surprisal.LM contribute significantly to improving a the prediction of the amplitude of the N400 by a linear model that only includes the baseline (adding SU.SGM: χ^2^ = 74.51, *p <* 0.0001, Δ*AIC* = 70.5; adding SU.SGM: χ^2^ = 89.67, *p <* 0.0001, Δ*AIC* = 85.7).

**Tab. 4:**
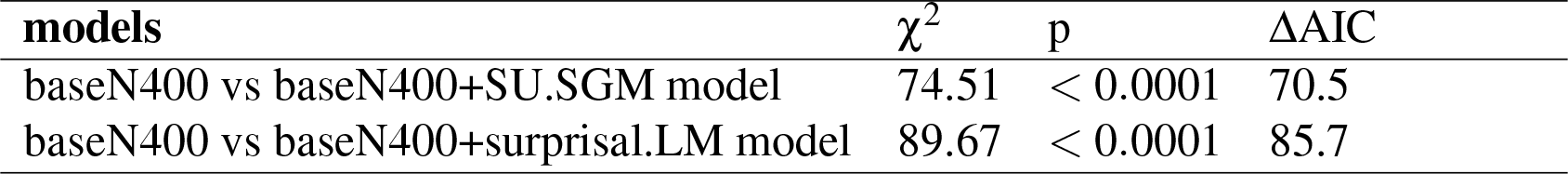
Results of analyses of variance and AIC reduction between a linear model of the N400 fit with only its N400 baseline and models fit in addition with either with SU.SGM or surprisal.LM.

Furthermore, we measured the improvement in N400 prediction by incorporating SU.SGM into a model already fitted with surprisal, and vice versa. Once again, the models were fitted with per-subject random slopes and random intercepts, along with per-word random intercepts. We conducted a two log-likelihood test between the models reporting both χ^2^ and Δ*AIC*.

Table 5 contains the results of two separate tests. The top row shows the comparison between a linear mixed effect model fitting surprisal and a model fitting both surprisal and SU.SGM. Adding SU.SGM to a model fitted with surprisal significantly improves it (χ^2^ = 69.58, *p <* 0.001 with Δ*AIC* = 63.5). The bottom row instead compare a model fitting only SU.SGM to a model fitting both surprisal and SU.SGM, with results indicating a significance improvement after the introduction of surprisal (χ^2^ = 84.74, *p <* 0.001 with Δ*AIC* = 78.7).

**Tab. 5:**
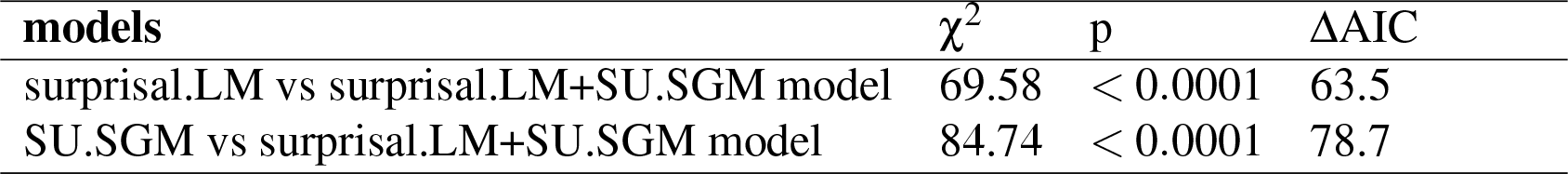
Results of analyses of variance and AIC reduction between a linear model of the N400 fit with lexical surprisal and a model fitted in addition with SU.SGM.

| **models** | $\chi^2$ | p | $\Delta AIC$ |
| --- | --- | --- | --- |
| surprisal.LM vs surprisal.LM+SU.SGM model | 69.58 | < 0.0001 | 63.5 |
| SU.SGM vs surprisal.LM+SU.SGM model | 84.74 | < 0.0001 | 78.7 |

Furthermore, in Supplementary Material C, we present a comparison of the effects on the N400 of the SU using untrained and trained models for both the SGM and LM. In addition, Supplementary Material D presents a comparison between SU.SGM and surprisal computed by GPT-2. Although this comparison is not our primary focus given that GPT-2 has a significantly larger number of parameters and a much larger training corpus compared to our SG model, we included the results in the Supplementary Materials for the sake of completeness.

### 3.3 Predicting the EEG time-course and topographical distribution

In order to delineate the relation between semantic update and surprisal with brain activity in more detail and beyond the N400, we conduct an exploratory analysis fitting a series of linear mixed effect models on each time-point of the EEG signal averaged across a region of interest defined over centro-parietal electrodes, the same as used for the N400 analyses reported in the previous sections and corresponding to the electrodes employed by Frank et al. (2015) to define the N400. The linear models followed the same structure as in Section 3.1, controlling for the baseline defined as the pre-word onset activity and with either SU.SGM or surprisal.LM as predictor of interest. We report the beta-coefficients over time (from 0 to 650 ms after word onset) relative to the estimates of these predictors of interest (Figure 6).

**Fig. 6:**
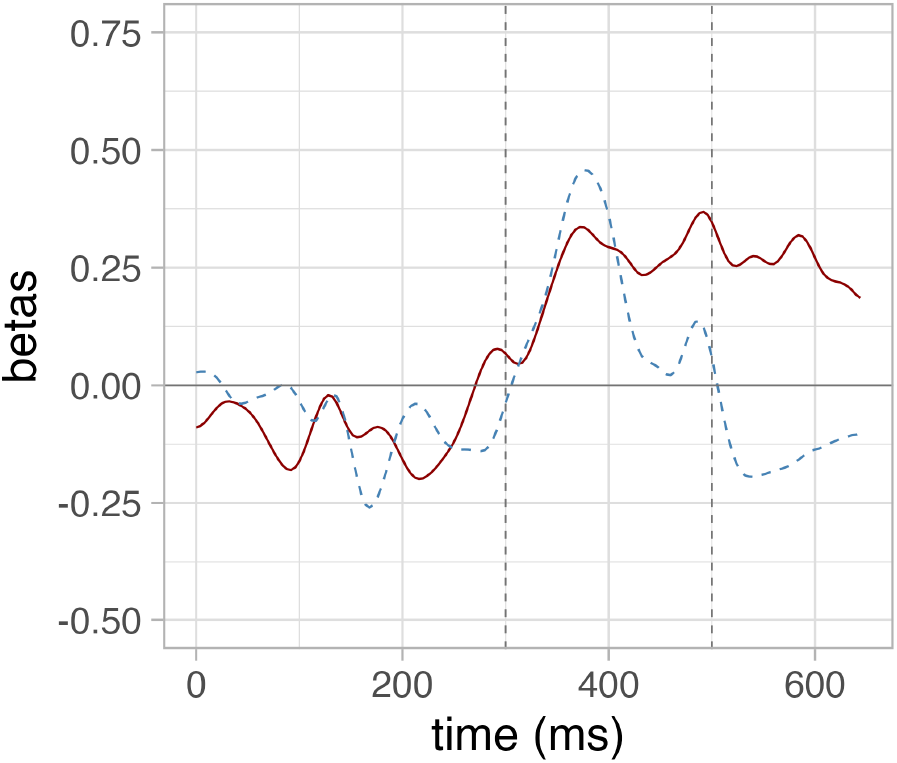
The time-wise results of a series of independent linear mixed effect models predicting the EEG signal in a ROI defined over N400-sensitive electrodes as a function of the SU.SGM (red) and surprisal.LM (dashed blue).

Furthermore, we replicated the same time-wise analyses on each separate electrode in the dataset and plotted the results in a series of topographical maps representing the distribution of the fit between SU.SGM or surprisal to electrophysiological activity using 50 ms wide non-overlapping time-windows. Figure 7 displays the topographical distribution the b-coefficients estimated by linear mixed effect models predicting EEG activity using SU.SGM or surprisal.LM respectively. Here it is evident again how the effect of SU.SGM extends the whole time-span of the N400 and beyond at centro-parietal electrode positions. Surprisal.LM shows a similar distribution, although limited to the early temporal phases of the latencies of the N400 ERP component. Moreover, both regressors display an inverse effects in frontal regions at later latencies. For SU.SGM this is mainly limited to the 650 to 700 ms time-window and only slightly left-lateralized, whereas for surprisal this phenomenon starts to appear right after the N400 time-window, around 500 ms post stimulus onset and is more strongly left-lateralized.

**Fig. 7:**
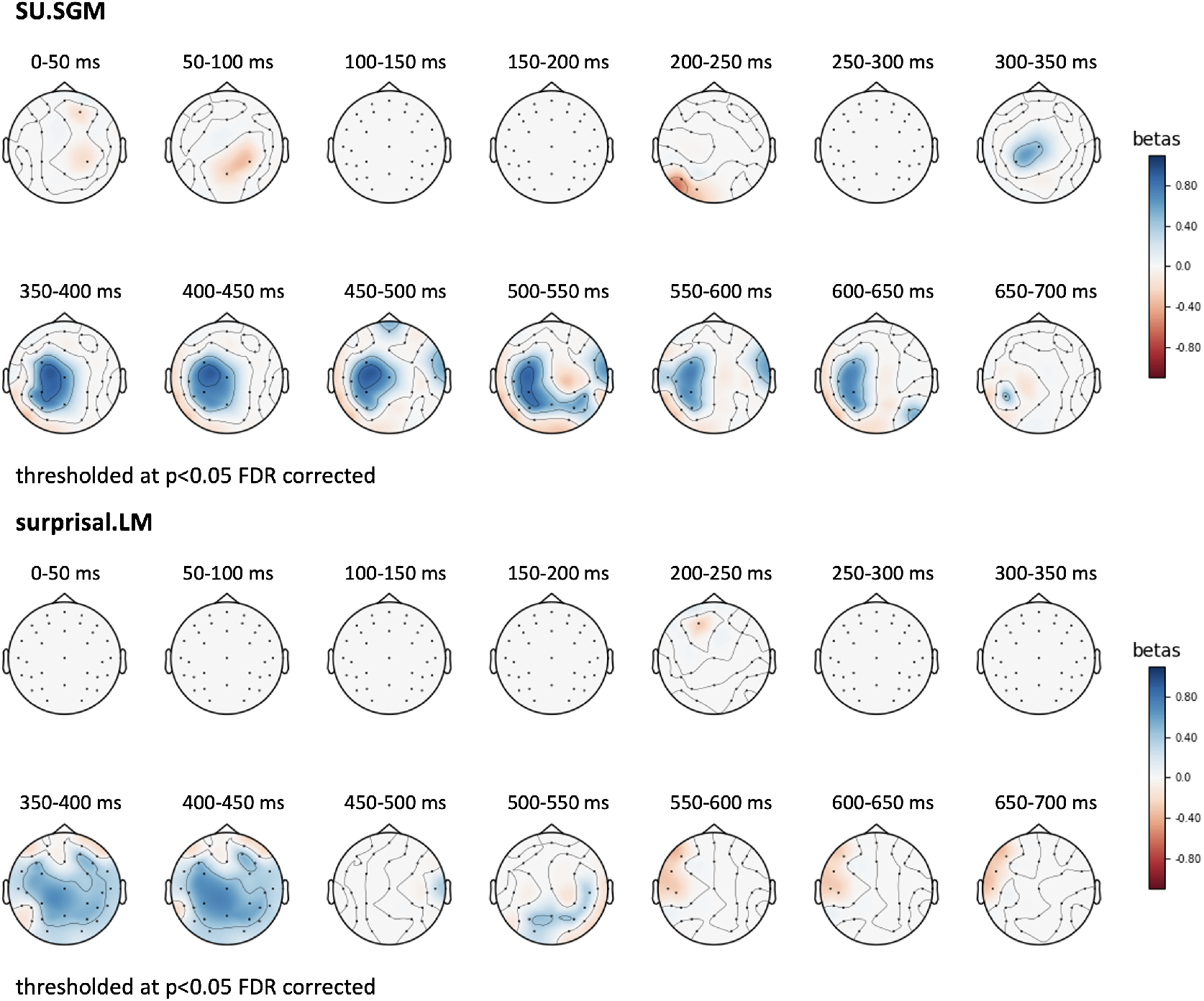
Topographical plot of the b-coefficients estimated by a linear mixed effect model predicting the EEG signal over time – from 0 to 600 ms post stimulus onset – as a function of **SU.SGM** (top) and **surprisal.LM** (bottom). We used 50 ms wide non-overlapping time-windows, and results are thresholded at p ¡ 0.05, FDR corrected.

Additionally, we conducted a latency analysis using the method introduced by Luck (2014) to determine the time point at which the peak effect (represented by the beta coefficients) of SU.SGM and surprisal.LM occurred over the centro-parietal electrodes. The method involves identifying a specific time point at which the area under the curve prior to that time point represents 50% of the total area under the curve for the entire time course.

First, we performed the analyses at the group level, using the beta coefficients obtained from the linear mixed-effects model reported in Figure 6. The results revealed a latency of 392 ms for surprisal.LM and 492 ms for SU.SGM and thus a numerical difference of 100 ms between the effects of surprisal and SU.SGM.

Next, we conducted the same analysis individually for each subject, utilizing a linear mixed-effects model that excluded the per-subject random slope and intercept. We subjected the latencies of both predictors to a t-test. Although the average latency of the SU.SGM effect was 397.33 ms, which was later than the latency of the surprisal.LM effect (373.33 ms), this difference was not statistically significant (*t* = 0.851, *p* = 0.404), possibly due to high inter-subject variability.

## 4 Discussion

Although there is a substantial amount of evidence linking the N400 to the processing of language, there is still a great deal of debate as to the nature of the processes it represents and the mechanisms that give rise to it. Here, following the results reported by Rabovsky et al. (2018), we advanced and tested the hypothesis that the N400 might signal the update of the implicit predictive representation of the meaning of a sentence during comprehension. This was accomplished by implementing the process in the form of a cognitively motivated artificial neural network model of predictive sentence comprehension (the Sentence Gestalt model) trained on a large language corpus, and by quantitatively mapping its internal dynamics (Semantic Update) to the N400 amplitude and more generally to the EEG measured during sentence comprehension. The usage of naturalistic training material allowed the models to learn from a linguistic environment as complex and close as possible to the one humans experience in the course of their lives. On a more technical level, it also allowed the models to be presented with the same stimuli used during human electrophysiological data collection, allowing for direct and quantitative comparison between model-derived variables and brain activity.

We quantitatively compared this account of the N400 to the alternative hypothesis that this signal instead reflects lexical surprisal and is hence driven by predictive processing at the level of lexical items. We did so by comparing the effect of the Semantic Update of the Sentence Gestalt model to surprisal estimated by a language model trained on the same corpus.

### 4.1 The time-course of Semantic Update

Our results show a significant relationship between Semantic Update and the amplitude of the N400 ERP component in a time window between 300 and 500 ms post stimulus (see Section 3.1). This is confirmed by time-resolved analyses (see Section 3.3), which show that SU.SGM has a significant effect on the EEG activity in line with the latency and the topographical distribution (Figure 7) of the N400. More precisely, the effect of the Semantic Update of the SG model (SU.SGM) is significant from 300 to 700 post-word onset. This goes beyond the usual N400 time window but is in line with studies reporting N400 effects extending beyond the typical N400 segment (Rabovsky et al., 2008). These observations are in line with the results reported by Rabovsky, Hansen, and McClelland (2016) and Rabovsky et al. (2018) which observed how the magnitude of the Semantic Update estimated by a SG model trained on a small-scale world behaves strikingly similar to the amplitudes of the N400 under experimental conditions manipulating semantic, probabilistic and positional variables, among others. Overall, the results support the hypothesis that the N400 component of the event-related brain potential might reflect the change induced by an incoming stimulus in an implicit predictive representation of meaning during sentence comprehension.

### 4.2 Lexical and sentence meaning prediction

Sections 3.2 and 3.3 show that also lexical surprisal significantly explains electrophysiological activity in the N400 time frame, and that both SU.SGM and surprisal have a significant effect on the amplitude of the N400 when fit together. The surprisal’s fit partially overlaps with the one of the SU.SGM, especially at early stages of activity. Moreover, the temporal profiles of these two predictors, as shown in Figure 6, appear to indicate that while both exhibit a rapid improvement in fit around 300 ms after word onset, surprisal only exhibits a single prominent peak around 350 ms, followed by a decline after 400 ms. In contrast, the effect of SU.SGM remains consistently strong throughout the entire N400 time window and beyond.

One possible explanation of these results is that the activity after 300 ms from the presentation of a word and the amplitude of the N400, in particular, do not reflect a single monolithic process, but rather the combined activity of several cognitive sub-processes which are related yet distinguishable. It is possible that language processing is supported by predictive processes across several levels of representation, such as at the level of individual words (lexical level) and sentence meaning. The existence of at least two distinct predictive sub-processes at the level of lexical and semantic processing might be also compatible with a hierarchy of predictive processes (K. J. Friston, 2005; K. Friston & Kiebel, 2009). Specifically, it has been proposed that predictions and prediction errors are passed up the hierarchy from lower to higher processing stages. This might be reflected by initial parts of the N400 potentially signaling primarily lexical prediction errors (explaining the high peak for surprisal in the initial part of the N400), while later parts of the N400 may reflect primarily sentence meaning related prediction errors (explaining the continued high correspondence with SU.SGM). This also seems in line with Nieuwland et al. (2020)’s analysis which revealed temporally distinct (though overlapping) effects of predictability and plausibility on the N400. In particular, lexical predictability negatively correlated with widespread activity peaking around 350 ms after word onset. By contrast, plausibility was associated with a smaller, right-lateralized effect that started after the peak of the lexical predictability and continued after the end of the N400 time frame. In light of these results, the authors propose that semantic facilitation of predictable words is produced by a cascade of processes that activate and integrate contextual word meaning into semantic representations at the level of the sentence.

Theories of predictive and parallel processing have also found support from several additional brain-imaging studies. Using language models trained on different levels of linguistic description, Lopopolo et al. (2017) showed that predictive processes may be decomposed in separate domain-specific sub-processes mapped onto distinct cortical areas. This hypothesis is supported also by Heilbron, Armeni, Schoffelen, Hagoort, and de Lange (2020) who observed dissociable EEG and MEG neural signatures of lexical, syntactic, phonemic, and semantic predictions.

SU.SGM and surprisal are measures that presumably represent two different processes, and they tackle these processes from two different levels of analysis framed along the lines of Marr’s distinction between the *computational* and *algorithmic* (Marr, 1982) levels. As noted in the introduction, the computational level is concerned with the goal of the process carried out by a system. The algorithmic level, on the other hand, refers to the representations and operations involved in the process. Semantic Update measures the change over time of the SG model’s internal hidden layer activity during sentence processing. Therefore it is concerned with the representations and operations (for instance the recursive integration over time implemented by the SG layer) used by the system to perform the task of sentence comprehension. It thus offers a description of the algorithmic level of the process. For this reason, the observed effect of SU.SGM on the amplitude of the N400 might reflect the internal dynamics of a meaning construction mechanism signaled by the component. Surprisal, on the other hand, is obtained from the probability assigned by a language processing system to an incoming word given its context of utterance. Surprisal quantifies how much the system is off with regard to the estimation of the probability of that word appearing and it can be interpreted as lexical prediction error. Having this in mind, it is therefore a measure of the performance of a system in solving the task of predicting the next word, therefore making it a descriptor of the computational level of the process, in the sense in which Marr defines it.

### 4.3 The SG model and different interpretations of surprisal

In the present study, we have adopted an interpretation of word surprisal as lexical prediction error and contrasted it with semantic update, a measure of how the implicit distributional representation of sentence meaning changes during sentence processing. An alternative interpretation, presented by Levy (2008), suggests that lexical surprisal can instead be understood as the change in the probability of structural interpretations caused by incoming words. Thus, according to this theory, the relationship between N400 amplitudes and surprisal would suggest that N400 amplitudes correspond to the amount of update or change of the structural interpretation of the sentence induced by the current word, which is somewhat related to Semantic Update in the SG model. However, while Levy’s formulation of sentence processing focuses on probability distributions over syntactic structures, the SG model estimates probabilities of sentence meanings in the sense of pairs of thematic roles and fillers involved in the described event.

In addition, we propose that the difference between the SG model’s Semantic Update and surprisal can be characterized in terms of function and mechanism. Levy (2008) focuses on the functional and probabilistic definition of sentence comprehension, which is the conditional probability between sentence and structure. On the other hand, our analyses of the SG model focus on the mechanism generating intermediate representations (high dimensional distributed pattern of activity) and the changes they undergo during processing, which we see as presumably reflecting the update of the neural representation of sentence meaning. Semantic Update measures the update of the distributed internal representation of the sentence driven by input words, rather than the update of the explicit probability of sentence interpretations. This internal representation is the product of a recursive integrative mechanism, which ultimately supports estimating the probability distribution. Thus, while Levy’s theory seems to address the computational level, the SG model addresses the algorithmic level of sentence processing.

Besides, it is important to note that both surprisal and the SG model’s Semantic Update have a significant effect on N400 amplitudes even when considered together as predictors and that the temporal and topographical distribution of their distinct effects does not entirely overlap, supporting the view that they are at least partially related to different processes.

### 4.4 Computational modeling: comparison of functions and internal representations

When comparing Semantic Update and surprisal, it’s important to note that they are estimated from structurally similar but functionally different models. The SG and LM models have the same architecture for input, recurrent, and post-recurrent layers, but differ in their output layers. Their structural differences are necessary to approximate their functions: lexical and sentence meaning prediction. At the same time, both models might develop meaningful representations at the input layer in the form of word-level embeddings and at the subsequent recursive hidden layer in the form of sentence level representations as far as these representations are useful for performing the models’ tasks. Sentence level meaning representations might well be useful in predicting the next word. As a first step in investigating this hypothesis, we conducted an analysis in Supplementary Material B to examine how the Semantic Update of the recurrent layer of the Language Model (SU.LM) affected the N400 amplitude. Interestingly, we found that the effect of SU.LM on the N400 appears not to reach significance, unlike the SU.SGM, surprisal.LM: β = 0.08, *t* = 2.16, *p* = 0.055 (FDR corrected); SU.SGM: β = 0.22, *t* = 8.36, *p <* 0.001 (FDR corrected). Furthermore, the analysis revealed that both surprisal.LM and SU.LM have a significant effect when both are included in the model (surprisal.LM: β = 0.17, *t* = 3.24, *p* = 0.004, SU.LM: β = 0.10, *t* = 2.98, *p <* 0.006 FDR corrected); please see Supplementary Material B for more details. These findings potentially indicate that the different training tasks yield distinct internal representations, and if further studies replicate these results, it suggests that the internal representations generated by the SG model training might align more closely with the internal representations formed during human sentence comprehension, as observed in the N400.

In general, the extent to which the internal representations generated by next word prediction models capture meaning processing dynamics presents an intriguing open question that calls for further dedicated investigations. The remarkable performance of language models in next word prediction tasks provides suggestive evidence of such emergence. In fact, research conducted by Lindborg and Rabovsky (2021) explored the relationship between internal activation updates in GPT-2 and N400 amplitudes to investigate the potential connection between these models’ internal representations and the comprehension of meaning.

### 4.5 On the computational modelling approach

As apparent from the above discussion, while we have control over the architecture and training regime of the models, and thus over the function the models are approximating, the internal implicit representations emerging within the models and thus the processes approximated by their internal activations are somewhat less transparent. While we believe our interpretations of the processes approximated by these internal dynamics and output measures are plausible (see also Rabovsky et al. (2018) for further discussion concerning the hypothesized internal processes within the SG model), it is important to acknowledge that these are open for debate. Relatedly, our modeling approach may have some limitations because model measures (e.g., Semantic Update or surprisal) may correlate with dependent measures (e.g., the N400) to varying degrees without providing irrefutable evidence that the mechanisms underlying the dependent variable correspond exactly to the mechanisms implemented in the models. However, we believe that this computationally explicit quantitative approach represents a significant step forward in moving beyond discussions of vague verbal concepts such as e.g., lexico-semantic retrieval and access, and also beyond qualitative simulations using models trained on toy corpora.

### 4.6 Relation to late positivities?

Even though the observed effect of SU.SGM extends beyond the typical N400 segment and into the P600 time window, we still interpret this centro-parietal effect as modulation of the N400 rather than the P600 component (a subsequent positivity over centro-parietal areas, also referred to as late posterior positivity). This is because the effect goes against the typical polarity of the P600 and is in line with the expected polarity of the N400. Specifically, less expected sentence continuations usually go along with more negativity during the N400 (i.e., a larger negative N400), but with more positivity during the P600 (i.e., a larger positive P600). In our study, we find that a larger Semantic Update (SU) in the SG model goes along with more centro-parietal negativity during the N400 time window and beyond the most typical N400 time window. This finding is directly opposite to any expected influence on the P600, which would be expected to show larger SU.SGM going along with more positivity during the P600 (Brouwer et al., 2017, 2021); also see next section for further discussion. It is worth noting that both components are temporally not restricted to their most typical time segments but may overlap in time (Chwilla, Brown, & Hagoort, 1995).

However, interestingly, besides predicting N400 amplitudes, i.e., more negative amplitudes at centro-parietal electrodes, surprisal appears to be have an inverse effect on activity recorded in anterior regions following the N400 time-window (Figure 7), indicating a potential relation between the processes of lexical prediction as instantiated by our predictor and a frontal post N400 positivity. This seems in line with previous evidence showing positive components in anterior areas after the N400 (Thornhill & Van Petten, 2012; Petten & Luka, 2012a; DeLong, Quante, & Kutas, 2014; Brothers, Wlotko, Warnke, & Kuperberg, 2020; DeLong & Kutas, 2020; Kuperberg, Brothers, & Wlotko, 2020). The exact functional interpretation of these late anterior positivities in language processing is still actively debated. Kuperberg et al. (2020) argue that the post N400 frontal positivity might reflect a successful update of the meaning representation as a function of novel unpredicted input, whereas Thornhill and Van Petten (2012) interpret it as a function of lexical predictability independent from semantic similarity. One picture that seems to be emerging is that different from the late posterior positivity (P600; see above and next section), the late anterior positivity is increased for sentence continuations that are unexpected yet not implausible. This would fit with the fact that the EEG dataset we used was recorded during the processing of naturalistic sentences that contained variations of expectancy as encountered in everyday life, but did not include semantic violations.

### 4.7 Comparison to alternative cognitive models of the N400

Our results seem to indicate that the N400 might be indexing the change induced by an incoming stimulus to the representation of sentence meaning as recorded from the recurrent layer of a SG model, and that larger sentence meaning update continues to predict more negative ERP amplitudes over centro-parietal regions beyond the typical N400 segment. These conclusions are seemingly at odds with Brouwer et al. (2017) and Brouwer et al. (2021) that claim that the N400 instead indexes lexical retrieval, i.e. the activation from semantic memory of the meaning of each separate word in a sentence during its processing, and that the P600, instead, correlates with integrative processes aimed at constructing sentence level semantic representations. It is important to note that our results seem compatible with both word and sentence level contributions to the amplitude of the N400, which seem to partially overlap in time, a finding that seems at least partially consistent with Brouwer’s view on the N400 - lexical surprisal does not exactly correspond to Brouwer et al’s measure of the N400 because it is based on word form rather than word meaning, but our current study cannot disentangle both measures (however, please also note the potential alternative interpretation of surprisal as referring to sentence interpretation update as suggested by Levy, 2008, discussed above). Most importantly, however, our results provide strong evidence against the view that the P600 reflects sentence meaning update. If the P600 would reflect sentence meaning update during naturalistic reading as suggested by Brouwer et al. (2017, 2021), one would expect that larger SU.SGM (which reflects sentence meaning update) would predict more positive amplitudes at centro-parietal electrode sites in the P600 time segment starting at about 500 ms and extending beyond the end of Frank et al. (2015)’s recording segment. Instead, as noted above, what we observe is that a larger SU.SGM continues to predict more negative amplitudes at centro-parietal electrodes beyond the typical N400 window and into the P600 time window.^3^

How can we explain this discrepancy that Brouwer et al. (2017, 2021) reported that larger P600 amplitudes could be explained by larger sentence meaning update in their small-scale model, while we find in our quantitative analysis during naturalistic reading that larger sentence meaning update predicts more negative centroparietal ERPs during the P600 segment? It is important to note that both of Brouwer’s simulations concerned very specific experimental conditions. The 2017 model focused on reversal anomalies, i.e. sentences such as e.g., “Every morning at breakfast, the eggs would eat…” (Kuperberg, Sitnikova, Caplan, & Holcomb, 2003). In the model, the small N400 observed in these situations is taken to reflect facilitated lexical access due to semantic priming (e.g., from breakfast and eggs to eat) and the large P600 is taken to reflect more effortful sentence meaning update, due to the incongruency that the eggs in the sentence are agents of an eating action even though they are not usually able to do so. An alternative explanation of the same pattern of results is that the small N400 reflects an initial *semantic illusion*, in which the readers temporarily take the eggs to be the patient rather than the agent of the eating action, based on semantic priors (Kim & Osterhout, 2005; Rabovsky et al., 2018) and in line with good enough approaches to language comprehension (Ferreira, Bailey, & Ferraro, 2002; Ferreira, 2003). In this view, the large P600 might reflect an internal revision, i.e., the process that after the initial illusion participants detect that there is an anomaly in the sentence and revise their internal sentence representation so that it matches the syntactic structure. Thus, even though the large P600 can be interpreted and modeled as large sentence meaning update (as done by Brouwer et al. (2017)), it is arguably a very specific update process potentially including an internal revision process. The interpretation of the P600 as reflecting an internal revision process is compatible with long held views on the P600 (even though this interpretation has initially focused on the P600 as reflecting syntactic revision processes (Kaan & Swaab, 2003; Friederici, Mecklinger, Spencer, Steinhauer, & Donchin, 2001), it has been expanded to revisions of internal representations more generally (Petten & Luka, 2012b; Rabovsky & McClelland, 2020)). The second simulation of an N400 and P600 pattern (reported in Brouwer et al. (2021)) follows the same lines of the one above in that the observed increased P600 might again reflect an internal revision process rather than a simple sentence meaning update. Specifically, the experimental conditions were as follows (based on a study conducted by Delogu, Brouwer, and Crocker (2019)):

|  |  |
| --- | --- |
| Baseline: | John entered the restaurant. Before long, he opened the menu. |
| Event-related violation: | John left the restaurant. Before long, he opened the menu. |
| Event-unrelated violation: | John entered the apartment. Before long, he opened the menu. |

The event-related violation was the only condition that elicited a large P600. Since aspects of the context (e.g., “the restaurant”) and event (”opened the menu”) appear to be related, it is possible that, upon first reading, participants may initially believe that the sentence makes sense, resulting in a reduced N400, before realizing that something is off and revising their initial interpretation (resulting in a larger P600). Alternatively, as proposed by Brouwer et al. (2021), the small N400 can be explained as reflecting facilitated lexical access due to priming, while the P600 can, instead, reflect sentence meaning update. However, according to this latter explanation, one would expect an equally large P600 for the event-unrelated violation in which the sentence meaning update (in the absence of illusions and revisions) would be equally large. Crucially, the data did not show this: The P600 in the event-unrelated violation was equally small as in the baseline condition (Delogu et al., 2019). This pattern of results (of a large P600 only in the event related violation condition) makes sense from the perspective that the P600 reflects revision processes as these revision processes would be expected in the event-related violation but not in the event unrelated violation.^4^

To sum up, in both experimental settings in which Brouwer et al. (2017, 2021) simulated larger P600 amplitudes as larger sentence meaning update, the increased P600 might be alternatively explained as reflecting internal revision processes. Against this backdrop, our finding that larger sentence meaning update as measured by a large scale SG model during naturalistic reading does definitely not predict larger P600 amplitudes (instead predicting more negative amplitudes at centro-parietal electrode sites beyond the N400 time segment) might be seen as further evidence strengthening the view that the increased P600 observed in the specific experimental conditions targeted by Brouwer et al. (2017, 2021)’s models might rather reflect internal revision processes and that under natural conditions larger sentence meaning update is reflected in larger N400 amplitudes.

Our finding that SU.SGM explains N400 amplitudes as the average activity in the 300-500 ms time segment even when surprisal is accounted for (see Section 3.2) seems to provide evidence against the model by Fitz and Chang (2019) who link the N400 to purely lexical prediction error. However, our results seem consistent with lexical contributions to the earlier part of the N400 (see Sections 3.2 and 3.3), which thus seems partially consistent with their claims (please note that in the current study we do not differentiate between lexical contributions at the level of word form versus word meaning). Our results do not seem to speak to Fitz and Chang (2019)’s account of the P600 as a sequencing prediction error. However, the results of reversal anomalies with two animate event participants, such as “The fox hunted the poacher”, which do not include sequencing prediction errors but produce large P600 amplitudes (van Herten, Kolk, & Chwilla, 2005), seem inconsistent with their implementation.

## 5 Conclusions

In the present study, we quantitatively investigated the hypothesis that mechanisms underpinning the N400 ERP component might be related to the update of an implicit predictive representation of sentence meaning during language comprehension, and quantitatively compared it to the alternative hypothesis that the component is instead signaling lexical prediction error. These quantitative investigations were afforded by training an artificial neural network model of sentence comprehension, called the Sentence Gestalt (SG) model (McClelland et al., 1989) on a large scale naturalistic corpus, and directly comparing the update of the distributed predictive representation of sentence meaning after each incoming word (Semantic Update, or SU.SGM) with the EEG recordings obtained during sentence reading. Our work corroborates and crucially extends previous studies which have shown that the Semantic Update of an SG model trained on a small synthetic environment responds similarly to the N400 amplitude to a series of lexical semantic manipulations, including semantic congruity, cloze probability, semantic and associative priming, and repetition, among others (Rabovsky et al., 2018). The small scale training prevented a direct assessment of the similarity between electrophysiological data and the model’s dynamics as presented in the current study. In addition, our approach allowed to quantitatively compare the effects of the SG model’s update on the electrophysiological data with the effect of lexical surprisal estimated by a language model trained on the same language corpus.

The analyses reported in this paper demonstrated that there is a significant relationship between the amplitude of the N400 component and both the SU.SGM recorded from the SG model (Table 1) and surprisal from a LM (Table 2), with surprisal predicting especially the early part of the N400 and SU.SGM predicting also the later part of the N400 time window. In addition, we observed that the effect of SU.SGM on N400 amplitudes remains significant after controlling for surprisal, and vice versa. This seems to indicate that the activity corresponding to the N400 component might not correspond to a single monolithic process but might rather reflect at least two distinct sub-processes at the level of lexical and sentence meaning prediction error.

## Acknowledgements

The research has been funded by the Deutsche Forschungsgemeinschaft (DFG)- Project-ID 318763901 - SFB1294 (project B09 to Milena Rabovsky and Sebastian Reich), and the Emmy Noether grant RA 2715/2-1 to Milena Rabovsky.

## Footnotes

1 The decision to use pre-trained FastText embeddings for training the SG model is based on both theoretical and computational factors. The use of the embeddings is limited to the training of the SG model and allows us to implement the theoretical distinction between predicting sentence meaning and lexical item prediction. Thus, it is important to note that FastText embeddings are not part of the SG model’s architecture but are used as semantic representations of the training data necessary to approximate the semantic nature of the SG model’s function. Using embeddings maintains the SG model’s structure and training task consistent with its original formulation, which used manually coded binary semantic feature representations (McClelland et al., 1989). Embeddings enable the representation of a broader range of words necessary for the large-scale corpus training procedure adopted in our study.

2 To illustrate, imagine if we already knew all the factors that affect N400 amplitudes. Controlling for all these factors would remove the variance we seek to explain. Our theory was developed to account for all the variance induced by factors known to influence the N400 (see Rabovsky et al. (2018), for further discussion). If our cognitive theory failed to capture the influence of all the factors modulating the N400, it would imply a flaw in our theory. Therefore, in our view, controlling for factors known to influence the N400 is inconsistent with our theoretical objective to provide a theory that explains the natural variation of N400 amplitudes across sentences, encompassing all the variation induced by factors known to influence the N400.

3 In the present study, we predict the EEG data from the negative of SU, in order to accommodate the fact that the N400 is a negative component, and the fact that we expect (and find) larger SU.SGM to predict more negative N400 amplitudes.

4 Brouwer et al. (2021) claimed that there was no possible explanation for this specific pattern of results and subtracted the complete activity in the N400 time segment from the activity in the P600 segment to correct for “component overlap”. Even though a partial component overlap between the N400 and P600 might sometimes be an issue, in this specific dataset the wave-forms from the three conditions had already rejoined after the N400 and before the P600 time segment (see Figure 1 in Delogu et al. (2019)). Because there was a large N400 in the event-unrelated violation condition, subtracting this negativity from the P600 time segment indeed resulted in a large P600. Only after this major change of the data, the data fit the Brouwer model’s predictions.

