## Supplementary Materials for "Tracking lexical and semantic prediction error underlying the N400 using artificial neural network models of sentence processing"

#### A N400-sensitive electrodes

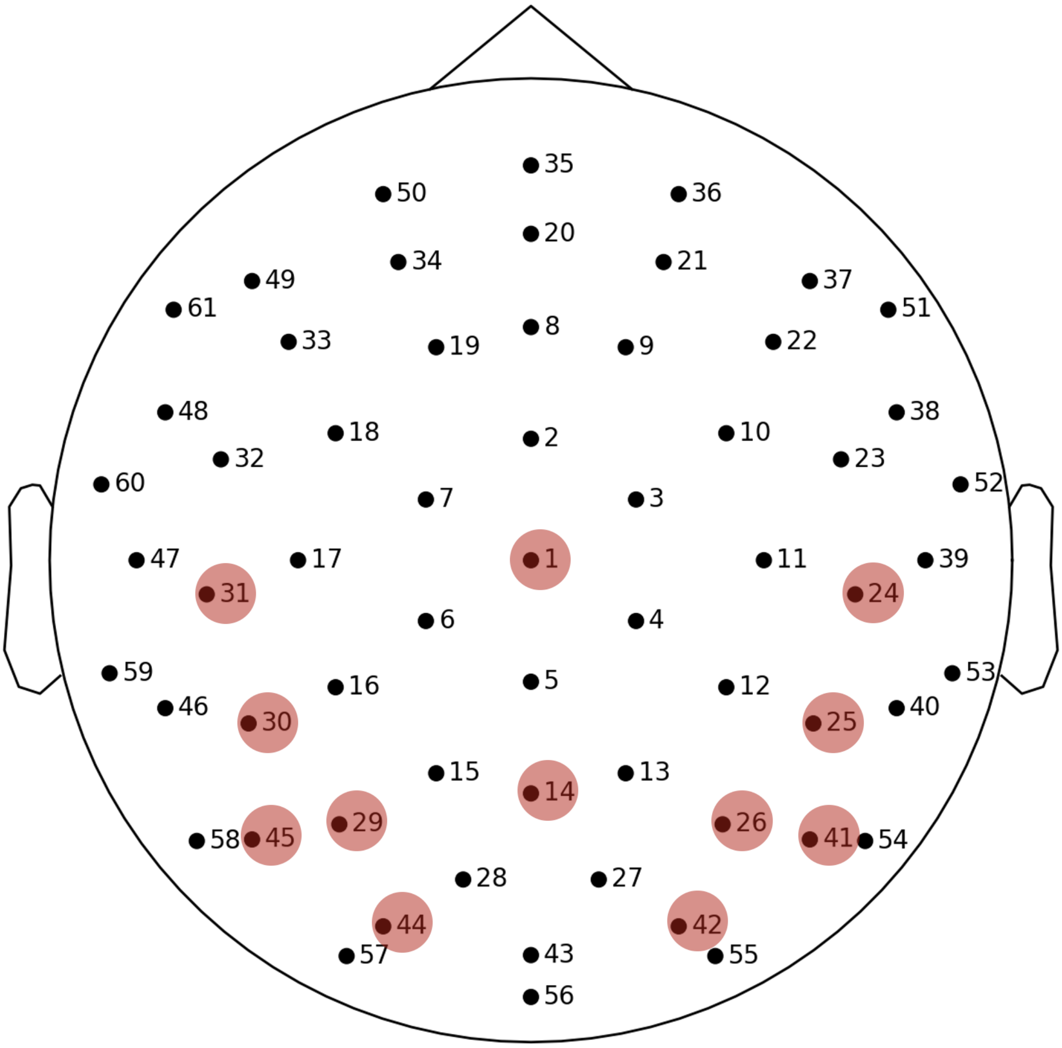


Fig. 8: Easycap M10 sensor array as used for the collection of the EEG data used in this study (Frank et al., 2015). The sensors highlighted in red are the one included in the definition of the N400 as used in the analyses based on the cluster used by Frank et al. (2015).

### B The Semantic Update of the Language Model

Here we assess the fit of the Semantic Update estimated from the recurrent layer of the LM (SU.LM) on the N400. This parallels the analyses conducted in the main text using SU.SGM, and as in those analyses we compare the effects of this regressor to surprisal estimated from the same LM (Section 3.2).

| | β | t | p |
|---|---|---|---|
| N400base | -0.41 | -20.94 | < 0.0001 |
| SU.LM | 0.08 | 2.16 | 0.055 |

Tab. 6: Linear mixed effect model fitted with the update of the LM (SU.LM) and aimed at predicting the amplitude of the N400 component.

| | β | t | p |
|---|---|---|---|
| N400base | -0.41 | -21.00 | < 0.0001 |
| SU.LM | 0.10 | 2.98 | 0.006 |
| surprisal.LM | 0.17 | 3.24 | 0.004 |

Tab. 7: Results of a model fitted with both surprisal and SU.LM and aimed at predicting the amplitude of the N400 component.

Table 6 shows the results of a linear mixed effect model predicting the N400 as a function of SU.LM. The model included also the N400 baseline and was fit with per subject random slope and intercept and per word random intercept. The effect of the SU.LM does not reach significance after correction ($\beta = 0.08$, $t = 2.16$, $p = 0.055$ FDR corrected). Table 7 contains the results of a linear mixed effect model fitted with SU.LM and surprisal.LM showing that both predictors derived from the language model are significant (surprisal: $\beta = 0.17$, $t = 3.24$, $p = 0.004$; SU.LM: $\beta = 0.10$, $t = 2.98$, $p = 0.006$ FDR corrected).

We assessed the improvement in the prediction of the N400 by incorporating SU.LM into a model fitted only with the N400 baseline. The models were fitted with per-subject random slopes and random intercepts, along with per-word random intercepts. We conducted a two log-likelihood test between the models reporting both $\chi^2$ and $\Delta AIC$. The same analysis concerning the effect of surprisal.LM are reported in Table 4 in the main text.

| **model** | $\chi^2$ | p | ΔAIC |
|---|---|---|---|
| baseN400 vs baseN400+SU.LM model | 32.18 | < 0.0001 | 28.2 |

Tab. 8: Results of analyses of variance and AIC reduction between a linear model of the N400 fit with only its EEG baseline and a model fit in addition with SU.LM.

The results reported in Table 8 indicates that the addition of SU.LM contributes significantly to improving a the prediction of the amplitude of the N400 by a linear model that only includes the baseline ($\chi^2 = 32.18$, $p < 0.0001$, $\Delta AIC = 28.2$).

We also measured the improvement in N400 prediction by incorporating SU.LM into a model already fitted with surprisal.LM, and vice versa. The models contains also the N400 baseline and were fitted with per-subject random slopes and random intercepts, along with per-word random intercepts. We conducted log-likelihood tests between the models reporting both $\chi^2$ and $\Delta AIC$.

| **models** | $\chi^2$ | p | ΔAIC |
|---|---|---|---|
| surprisal.LM and surprisal.LM+SU.LM model | 24.23 | $< 0.0001$ | 18.2 |

Tab. 9: Results of analyses of variance and AIC reduction between a linear model of the N400 fit with lexical surprisal and a model fitted in addition with SU.LM.

In Table 9 we report the results of analyses of variance and AIC reduction between a linear model of the N400 fit with surprisal.LM and a model fitted with both surprisal.LM in addition with SU.LM ($\chi^2 = 24.23$, $p < 0.0001$, $\Delta AIC = 18.2$).

In line with the main analyses, we replicated the same time-wise analyses on each separate electrode in the dataset and plotted the results in a series of topographical maps representing the distribution of the fit between SU.LM or surprisal.LM to electrophysiological activity using 50 ms wide non-overlapping time-windows.

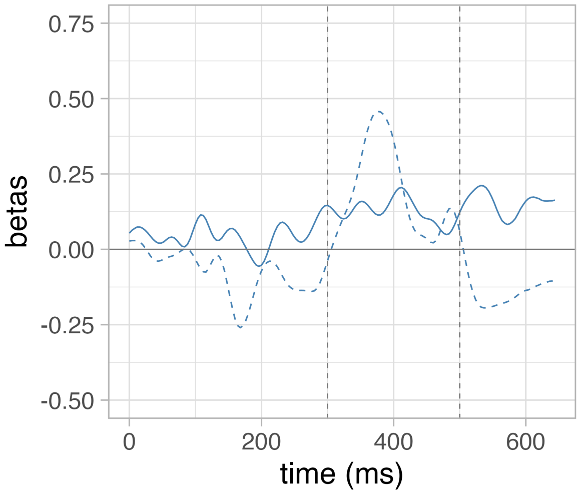


Fig. 9: The time-wise results of a series of independent linear mixed effect models predicting the EEG signal in a ROI defined over N400-sensitive electrodes as a function of the SU.LM (dark blue) and surprisal.LM (dashed blue).

Figure 9 compares the time course of the effect of SU.LM (dark blue) and surprisal.LM (dashed blue) on the activity recorded in N400 sensitive electrodes as defined by (Frank et al.,

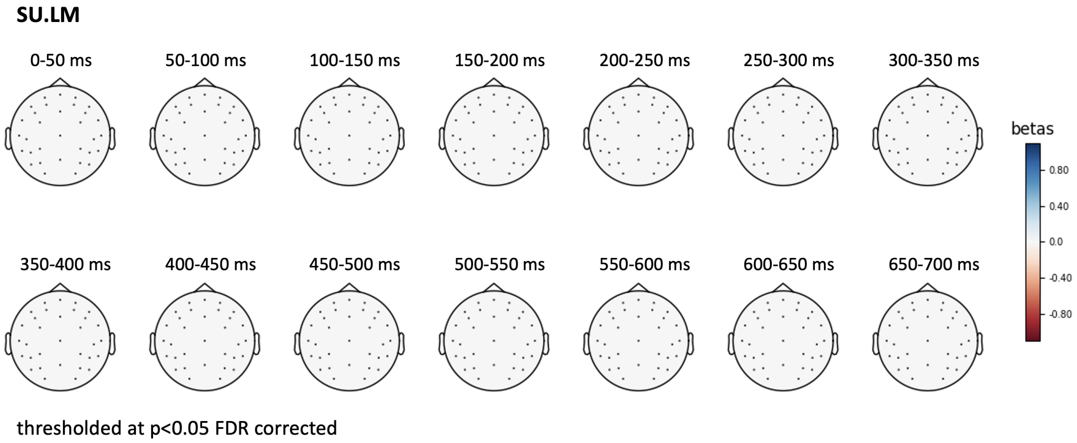


Fig. 10: Topographical plot of the β-coefficients estimated by a linear mixed effect model predicting the EEG signal over time – from 0 to 700 ms post stimulus onset – as a function of **SU.LM** over 50 ms wide non-overlapping time-windows (thresholded at p ¡ 0.05, FDR corrected).

2015). Figure 10 displays the topographical distribution the β-coefficients estimated by linear mixed effect models predicting EEG activity using SU.LM.

### C The effect of training

As humans acquire semantic knowledge about the world by experiencing their environment, the model learns knowledge about event semantics by being exposed to the statistic regularities in its environment – in this case the sentences contained in the RW-EN corpus – during training. In order to investigate whether training affects the similarity between the model's internal dynamics and the N400 amplitudes, we conducted similar analyses as in Section 3.1 using SU.SGM measures obtained from the same SG model before and after training.

Table 10 shows the results of a linear mixed effect models using as predictors of the N400 the SU.SGM computed from an untrained or a trained SG model. The effect of the SU.SGM of the untrained SG model is not significant ($\beta = -0.04$, $t = -1.63$, $p = 0.16$, FDR corrected). Additionally, we quantified the contribution of SU.SGM to the amplitude of the N400 before and after training by conducting two separate ANOVAs. The first ANOVA compared a linear mixed effect model that included only the N400 baseline with another model that also incorporated the SU.SGM from a trained SG model. The second ANOVA compared the same base linear mixed effect model with another model that included the SU.SGM from an untrained SG model. Table 11 reports the results of these comparisons. Only the SU.SGM obtained from the trained model appears to improve the fit to the amplitude of the N400 ($\chi^2 = 74.51$, $p < 0.0001$, $\Delta AIC = 70.5$), whereas the one from an untrained model does not ($\chi^2 = 3.61$, $p = 0.16$, $\Delta AIC = -0.4$).

| | untrained | | | trained | | |
|---|---|---|---|---|---|---|
| | β | t | p | β | t | p |
| N400base | -0.09 | -20.88 | < 0.001 | -0.41 | -21.01 | < 0.0001 |
| SU.SGM | -0.04 | -1.63 | 0.16 | 0.21 | 6.36 | < 0.0001 |

Tab. 10: Comparison of the effects of the SU.LM estimated from an untrained (left) and trained (right) SG model.

| **models** | $\chi^2$ | p | ΔAIC |
|---|---|---|---|
| baseN400 vs. baseN400+untrained SU.SGM | 3.61 | 0.16 | -0.4 |
| baseN400 vs. baseN400+trained SU.SGM | 74.51 | < 0.0001 | 70.5 |

Tab. 11: Results of an analysis of variance and AIC reduction between a linear model of the N400 without SU.SGM and one with SU.SGM from an untrained SG model (top) and between the same base model and one containing the SU.SGM values from a trained SG model (bottom).

The effect of LM training on the relationship between surprisal and the amplitude of the N400 was evaluated using the same approach. Table 12 displays the outcomes of two linear mixed effect models that predicted the N400 using surprisal from either an untrained or a trained LM. The effect of surprisal from the untrained LM was not found to be significant

($\beta = -0.01, t = -0.50, p = 0.62$, FDR corrected). ANOVAs were also employed to assess the impact of surprisal on the N400 amplitude before and after training. The first ANOVA compared a linear mixed effect model that included only the N400 baseline with another model that incorporated the SU.SGM from a trained SG model. Similarly, the second ANOVA compared the base linear mixed effect model with another model that included surprisal from an untrained LM. The results of these comparisons are presented in Table 13. It is evident that only the surprisal derived from the trained model improved the fit to the N400 amplitude ($\chi^2 = 89.67$, $p < 0.0001$, $\Delta AIC = 85.7$), whereas the surprisal from the untrained model did not yield significant improvements ($\chi^2 = 0.24$, $p = 0.88$, $\Delta AIC = -3.8$).

| | untrained | | | trained | | |
|---|---|---|---|---|---|---|
| | $\beta$ | t | p | $\beta$ | t | p |
| N400base | -0.41 | -20.86 | $< 0.001$ | -0.41 | -21.02 | $< 0.0001$ |
| surprisal.LM | -0.01 | -0.50 | 0.62 | 0.15 | 2.79 | 0.01 |

Tab. 12: Comparison of the effects of surprisal.LM estimated from an untrained (left) and trained (right) LM.

| **model** | $\chi^2$ | p | ΔAIC |
|---|---|---|---|
| baseN400 vs. baseN400+untrained surprisal.LM | 0.24 | 0.88 | -3.8 |
| baseN400 vs. baseN400+trained surprisal.LM | 89.67 | $< 0.0001$ | 85.7 |

Tab. 13: Results of an analysis of variance and AIC reduction between a linear model of the N400 without surprisal.LM and one with surprisal.LM from an untrained LM (top) and between the same base model and one containing the surprisal.LM values from a trained LM (bottom).

These results seem to indicate that the models' ability to predict the N400 amplitude is strongly affected by the exposure to their training environment. It is evident from these results that the SU.SGM and surprisal obtained from a trained models better approximates the N400 as compared to the ones obtained from random models, i.e. models with randomly initialized connection weights.

Recently, Schrimpf et al. (2021) analyzed the fit between fMRI and ECoG data in the frontal-temporal cortex and vectorial representations generated by 43 deep learning models. Similarly to our study, besides using fully trained models, they also evaluate the same models before training. Interestingly, they observed that untrained networks yield representations that still significantly predict fMRI data, although training significantly improves fit. These results led to the proposal that the architecture of the networks can work as brain models of language even without extensive training because the hierarchical structures implementing the deep learning networks might resemble similar neural mechanisms implemented by the cortical regions under analysis. We cannot conclude, based on our observations, that the architecture of the SG model alone – i.e. without training on a cognitively plausible task – is enough to approximate the electrophysiological processes under scrutiny. This partially diverges from Schrimpf et al. (2021)'s position. Nonetheless, we think it is important to stress that the difference between their and our conclusions might simply be due to the fact that the most successful models in their study were implemented using architectures – such as the multi-head attention mechanism (Vaswani et al., 2017) – which were not used for the present iteration of the SG model. Moreover, instead of analyzing fMRI and ECoG data, we focused on EEG activity and in particular on the amplitude of the N400. Therefore, it could also be the case that our results simply reiterate the fact that the N400 ERP component's behavior evolves during an individual's experience of the statistical regularities of their environment, paralleled by the activity of the Sentence Gestalt layer during training epochs. Importantly, our results also emphasize that the implicit semantic prediction error reflected in N400 amplitudes inherently depends on the statistics of the environment as the predictions formed by the model (and presumably by human comprehenders) are generated based on the experience of these statistics.

Previously, Rabovsky et al. (2018) also investigated the effect of training on the SG model's ability to simulate N400 amplitudes, specifically during the processing of sentences containing semantically incongruent nouns. They reported that the SU.SGM shows at first an increase and later a decrease with additional training. These results are in line with the variation of the N400 during human language acquisition, which also first increases and then decreases across development (Friedrich & Friederici, 2004; Atchley et al., 2006; Kutas & Iragui, 1998). Moreover, Rabovsky et al. (2018) observed that the output layer activation approximates more and more the probability distributions embodied in the training corpus. This second point is in line with our results in confirming the role of training in improving the fit between a computational model and human data, mediated by the fit to the statistics of the world.

### D Controlling for surprisal from large-scale transformer models

In this section, we assess the fit of Semantic Update on the N400 together with surprisal estimated from GPT-2. In the main text, we compared the SU.SGM to the surprisal estimated from a LM with a comparable architecture that was trained on the same corpus (Section 3.2). This was because we felt that the disparity in number of parameters and training data between the SGM and state-of-the-art language models (such as GPT-2) could act as a confound in the assessment of the performance with regard to the N400 amplitude and would prevent a fair comparison between the mechanisms and hypotheses implemented by the models.

We first fit a linear mixed effect model predicting the N400 as a function of both SU.SGM and surprisal estimated by GPT-2. The model included also the N400 base as described in the main text and per-subject random slopes and random intercepts, and per-word random intercepts.

| | β | t | p |
|---|---|---|---|
| N400base | -0.09 | -21.01 | $< 0.0001$ |
| SU.SGM | 0.10 | 3.80 | $< 0.001$ |
| surprisal.GPT-2 | 0.36 | 8.73 | $< 0.0001$ |

Tab. 14: Linear mixed effect model fitted with the SG model SU (SU.SGM) and surprisal estimated by GPT-2 and aimed at predicting the amplitude of the N400 component.

Table 14 contains the results indicating that even with the presence of surprisal.GTP-2 ($\beta = 0.36$, $t = 8.73$, $p < 0.0001$ FDR corrected), SU.SGM makes a significant contribution to the amplitude of the N400 ($\beta = 0.10$, $t = 3.80$, $p < 0.001$ FDR corrected), and vice versa.

We assessed the improvement in the prediction of the N400 by incorporating either the SU.SGM or surprisal.GPT-2 into a model fitted only with the N400 baseline. The models were fitted with per-subject random slopes and random intercepts, along with per-word random intercepts. We conducted a two log-likelihood test between the models reporting both $\chi^2$ and $\Delta AIC$.

| **models** | $\chi^2$ | p | ΔAIC |
|---|---|---|---|
| baseN400 vs baseN400+SU.SGM model | 74.51 | $< 0.0001$ | 70.5 |
| baseN400 vs baseN400+surprisal.GPT-2 model | 214.69 | $< 0.0001$ | 210.7 |

Tab. 15: Results of analyses of variance and AIC reduction between a linear model of the N400 fit with only its N400 baseline and models fit in addition with either with SU.SGM or surprisal.LM.

The results reported in Table 15 indicates that both the addition of SU.SGM and of surprisal.GPT-

2 contributes significantly to improving a the prediction of the amplitude of the N400 by a linear model that only includes the baseline.

We also measured the improvement in N400 prediction by incorporating SU.SGM into a model already fitted with surprisal estimated by GPT-2, and vice versa. The models were fitted with per-subject random slopes and random intercepts, along with per-word random intercepts. We conducted a two log-likelihood test between the models reporting both $\chi^2$ and $\Delta AIC$.

| **models** | $\chi^2$ | p | ΔAIC |
|---|---|---|---|
| surprisal.GPT-2 vs surprisal.GPT-2+SU.SGM model | 14.45 | $< 0.01$ | 8.4 |
| SU.SGM vs surprisal.GPT-2+SU.SGM model | 154.63 | $< 0.0001$ | 148.6 |

Tab. 16: Results of analyses of variance and AIC reduction between a linear model of the N400 fit with surprisal estimated by GPT-2 and a model fitted in addition with SU.SGM.

Table 16 contains the results of two separate tests. The top row shows the comparison between a linear mixed effect model fitting surprisal and a model fitting both surprisal.GPT-2 and SU.SGM. Adding SU.SGM to a model fitted with surprisal.GPT-2 significantly improves it ($\chi^2 = 14.45$, $p < 0.01$ with $\Delta AIC = 8.4$). The bottom row instead compare a model fitting only SU.SGM to a model fitting both surprisal.GPT-2 and SU.SGM, with results indicating a significance improvement after the introduction of surprisal ($\chi^2 = 154.63$, $p < 0.0001$ with $\Delta AIC = 148.6$).

Furthermore, we replicated the same time-wise analyses on each separate electrode in the dataset and plotted the results in a series of topographical maps representing the distribution of the fit between SU.SGM or surprisal to electrophysiological activity using 50 ms wide non-overlapping time-windows. Figure 11 compares the time course of the effect of SU.SGM (dark red) and GPT-2 surprisal (dashed green) on the activity recorded in N400 sensitive electrodes as defined by (Frank et al., 2015). Figure 12 displays the topographical distribution the β-coefficients estimated by linear mixed effect models predicting EEG activity using surprisal.GPT-2.

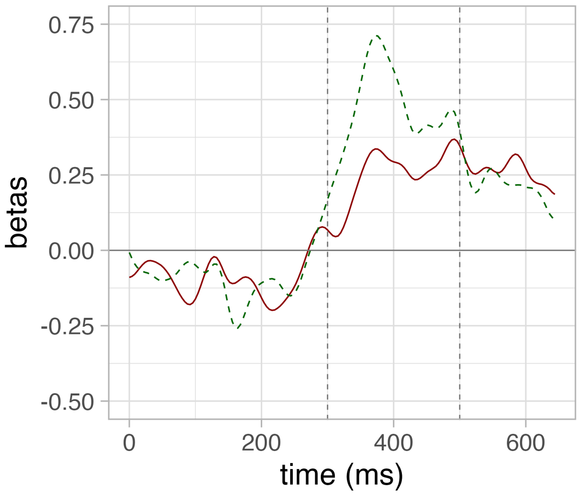


Fig. 11: The time-wise results of a series of independent linear mixed effect models predicting the EEG signal in a ROI defined over N400-sensitive electrodes as a function of the SU.SGM (red) and GPT-2 surprisal (green).

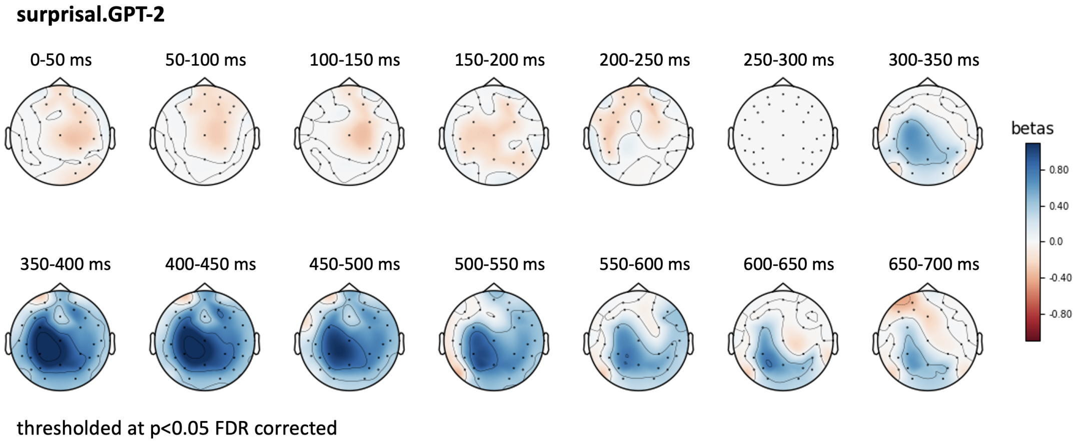


Fig. 12: Topographical plot of the goodness of fit of a linear mixed effect model predicting the EEG signal over time – from 0 to 700 ms post stimulus onset – as a function of **GPT-2 surprisal** over 50 ms wide non-overlapping time-windows (thresholded at p ¡ 0.05, FDR corrected).
